# Mechanism-based crosslinking probes capture *E. coli* ketosynthase FabB in conformationally-distinct catalytic states

**DOI:** 10.1101/2022.04.04.486996

**Authors:** Aochiu Chen, Jeffrey T. Mindrebo, Tony D. Davis, Woojoo E. Kim, Yohei Katsuyama, Ziran Jiang, Yasuo Ohnishi, Joseph P. Noel, Michael D. Burkart

**Affiliations:** Chemistry and Biochemistry, University of California San Diego, 9500 Gilman Drive, La Jolla, CA, 92093, USA; Jack H. Skirball Center for Chemical Biology and Proteomics, Salk Institute for Biological Studies, La Jolla, CA, 92037, USA; Department of Biotechnology, Graduate School of Agricultural and Life Sciences, The University of Tokyo, 1-1-1 Yayoi, Bunkyo-ku, Tokyo, 113-8657, Japan; Collaborative Research Institute for Innovative Microbiology, The University of Tokyo, 1-1-1 Yayoi, Bunkyo-ku, Tokyo, 113-8657, Japan

**Keywords:** ketosynthase, fatty acid synthase, decarboxylative condensation reaction, acyl carrier protein, protein crosslinked complex

## Abstract

Ketosynthases (KS) catalyse essential carbon-carbon bond forming reactions in fatty acid biosynthesis using a two-step, ping-pong reaction mechanism. In *E. coli*, there are two homodimeric elongating KSs, FabB and FabF, both of which possess overlapping substrate selectivity. However, FabB is essential for the biosynthesis of unsaturated fatty acids (UFAs) required for cell survival in the absence of exogenous UFAs. Additionally, FabB has reduced activity towards substrates longer than 14 carbons, whereas FabF efficiently catalyses the elongation of saturated C14 and unsaturated C16:1 acyl-acyl carrier protein (ACP) complexes. In this study, we solved two crosslinked crystal structures of FabB in complex with ACPs functionalized with long-chain fatty acid crosslinking probes that approximate catalytic steps. Both homodimeric structures possess asymmetric substrate binding pockets, suggestive of cooperative relationships between the two FabB monomers when engaged with C14 and C16 acyl chains. In addition, these structures capture an unusual rotamer of the active site gating residue, F392, potentially representative of the catalytic state prior to substrate release. These structures demonstrate the utility of mechanism-based crosslinking methods to capture and elucidate at near atomic resolution conformational transitions accompanying KS-mediated catalysis.

**Synopsis:** Crystal structures of KS-ACP crosslinked complex elucidate chain length preference and substrate processing mechanism of *E. coli* FabB.

## 1. Introduction

Fatty acid synthases (FASs) carry out a series of iterative biochemical transformations to produce fatty acids (FAs). Depending on their organization, FASs are classified as type I multidomain megasynthases, or type II multiprotein complexes comprised of discrete, monofunctional enzymes. Interestingly, Type II FASs, commonly found in bacteria and plant plastids, are considered the evolutionary progenitors of polyketide synthases that produce complex, biologically active molecules (Chen *et al*., 2018; Hillenmeyer *et al*., 2015). FASs iteratively extend and reduce two-carbon malonyl-CoA derived units. A ketosynthase (KS) begins each round of chain elongation by condensing a malonyl-CoA derived substrate with a growing fatty acid to generate a β-keto intermediate via a decarboxylative Claisen condensation reaction. Subsequently, a ketoreductase, a dehydratase, and an enoylreductase each catalyse three sequential reactions to fully reduce the β-ketone to a methylene. Central to the elongation cycle is the acyl carrier protein (ACP), which is required to shuttle thioester-tethered substrates to each enzyme active site (White *et al*., 2005). The thiol group of the ACP is derived from 4’-phosphopantetheine (PPant), which is post-translationally installed onto a conserved serine residue of the ACP. The protein-protein interactions (PPIs) of ACP and its partner enzymes are crucial for efficient substrate delivery, particularly in type II systems possessing discrete domains (Mindrebo *et al*., 2020c).

KSs catalyse the critical carbon-carbon bond forming reactions at the beginning of each FAS elongation cycle (Heil *et al*., 2019). The KS two-step, ping-pong mechanism consists of two half-reactions that include a transacylation and a condensation step. During the transacylation half-reaction, the acyl-chain from acyl-ACP is transferred to the catalytic cysteine of the *apo*-KS to generate an acyl-KS intermediate. During the condensation half-reaction, malonyl-ACP (mal-ACP) binds to the KS and undergoes decarboxylation to form a two carbon enolate ion that attacks the acyl-KS thioester to form β-ketoacyl-ACP. *Escherichia coli* possesses two elongating KSs, FabB and FabF, belonging to the ketoacyl-ACP-synthase I (KASI) and ketoacyl-ACP synthase II (KASII) families, respectively. The two KSs have partially overlapping function, but FabB specifically elongates *cis*-3-decenoyl acid (C10:1), the first committed unsaturated fatty acid (UFA) intermediate in *E. coli*. Strains lacking FabB are UFA auxotrophs, but those lacking FabF are viable without supplementation (Cronan *et al*., 1969; D’Agnolo *et al*., 1975; Feng & Cronan, 2009). In terms of the chain length selectivity, FabB has reduced activity towards substrates longer than 12 carbons whereas FabF retains robust activity on 14-carbon and 16-carbon FA substrates (Edwards *et al*., 1997).

Interactions between ACP and its partner enzymes (PEs) are transient, thus complicating the visualization of PE-ACP complexes that approximate key catalytic states. In recent years, the application of phosphopantetheine analogs outfitted with mechanism-based crosslinking probes mimicking fatty acyl substrates have advanced our understanding of the multistep KS reaction mechanisms and substrate specificities for both FAS and PKS (Chen *et al*., 2022; Milligan *et al*., 2019; Mindrebo *et al*., 2020b; Du *et al*., 2020; Mindrebo *et al*., 2021; Herbst *et al*., 2018). These probes contain selectively reactive warheads that form a covalent bond with the catalytic cysteine of KSs, resulting in the formation of a covalently trapped KS-ACP complex for structure elucidation using X-ray crystallography. Two classes of KS warheads, chloroacrylates and α-bromo carbonyl containing molecules (Worthington *et al*., 2006; Worthington *et al*., 2008), have been successfully applied in previous structural studies. Chloroacrylate crosslinked complexes, which tether the C_3_ position of the acyl chain to the KS active site via conjugate substitution, mimic an intermediate formed during the condensation half-reaction as demonstrated by previously published ACP-KS crosslinked structures of FAS and PKS (Milligan *et al*., 2019; Du *et al*., 2020; Mindrebo *et al*., 2021). The FabB-ACP α-bromo crosslinked complexes, which tether the C_2_ position of the acyl chain to the KS active site, mimic a transacylation intermediate. In contrast, FabF-ACP complexes trapped with α-bromo probes resemble an intermediate substrate delivery state typified by large conformational rearrangements of two KS active site loops (Mindrebo *et al*., 2020b; Mindrebo *et al*., 2021). The two loops, loop 1 and loop 2, form a double drawbridge-like gate (Gora *et al*., 2013) at the entrance of the active site pocket that regulate substrate processing. Current models suggest this gate is likely triggered by the interaction of loop 2 and acyl-ACP and the subsequent chain-flipping (Cronan, 2014) of ACP-sequestered cargo in the active site. The gate-open conformation has only been captured in crosslinked crystal structures, highlighting the utility of mechanism-based crosslinking probes.

In this study, we develop and apply two crosslinking probes bearing long-chain fatty acyl substrate mimetics to investigate the molecular basis of FabB substrate preference. The first probe, C14-chloroacrylate (C14Cl), mimics the final condensation reaction catalysed by FabB in the elongation cycle, while the second probe, C16:1-α-bromo (C16:1Br), represents the acylation reaction of FabB with a disfavoured substrate. Two crystal structures, FabB=C14-ACP and FabB=C16:1-ACP (where the ‘=’ sign denotes a crosslinked complex) were determined using C14Cl and C16:1Br as crosslinkers, respectively. Both structures possess a back gate comprised of a glutamine and a buried glutamate residue that can alternate between conformers to expand or limit the size of the fatty acyl binding pocket. The FabB=C14-ACP structure possesses a balanced substrate binding pocket arrangement that accommodates the C14 chain. However, the FabB=C16:1-ACP complex has distinct substrate binding pocket structures in each of the protomers. Importantly, the pocket conformations affect thioacyl coordination and alter the conformation of key catalytic residues related to KS gate function. Insights from these structures correlate well with FabB function and provide new insights into *E. coli* FAS and KS-directed metabolic engineering.

## 2. Materials and methods

### 2.1. Synthesis of crosslinking probes

C16:1Br, (Mindrebo *et al*., 2021) (*E*)-C3Cl, (Worthington *et al*., 2006), and (*E*)-C8Cl (Worthington *et al*., 2008) were prepared according to previous literature protocols. (*E*)-C14Cl and (*Z*)-C14Cl were prepared according to Scheme S1. See Supporting Information for complete synthetic details.

**Scheme S1:**
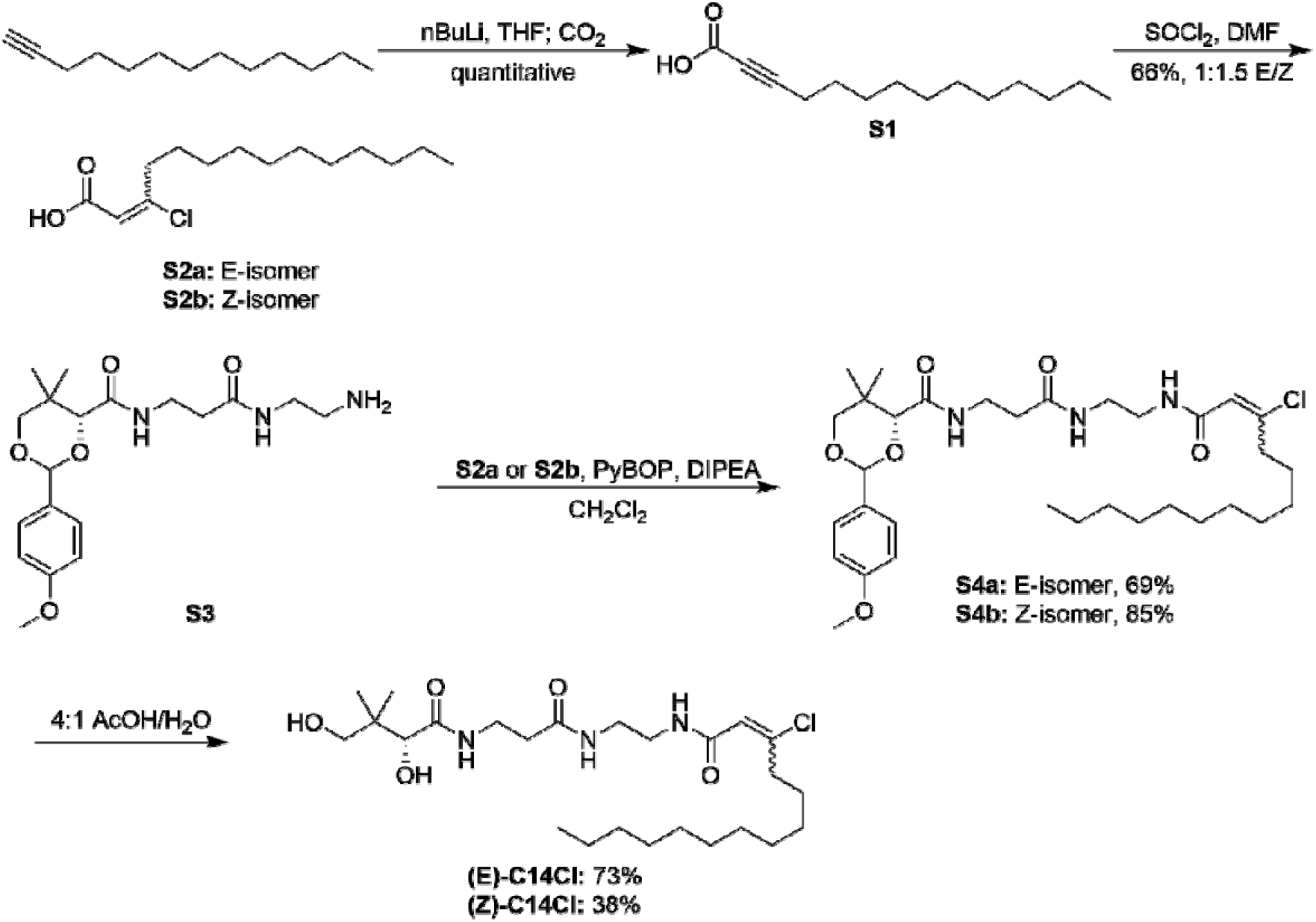
Synthesis of chloroacrylamide crosslinking probes. PyBOP= benzotriazol-1-yloxytripyrrolidinophosphonium hexafluorophosphate; DIPEA=*N*,*N*-diisopropylethylamine; AcOH=acetic acid.

### 2.2. Protein expression and purification

The N-terminal His-tagged FabB recombinant protein was expressed in *E. coli* BL21(DE3) and cultured in LB broth containing 50 mg/L kanamycin. Cells were grown at 37 °C until OD_600_ reached ~0.7 and induced by 0.5 mM IPTG for 4 h under the same temperature. Cell pellets were collected by centrifugation, lysed by sonication in lysis buffer (50 mM Tris pH 8.0, 150 mM NaCl, 10% glycerol), and underwent another centrifugation at 17,400 g for 1 h to clear the lysate. The His-tagged protein was pulled down by Ni-NTA resin (0.5 mL resin used per litre growth), washed with wash buffer (50 mM Tris pH 8.0, 150 mM NaCl, 10% glycerol) and 20 mM buffered imidazole, and eluted with 250 mM buffered imidazole. His-tag cleavage by bovine thrombin (2 U per 1 mg FabB) was performed while dialyzing against the dialysis buffer (50 mM Tris pH 8.0, 150 mM NaCl, 10% glycerol, 0.5 mM TCEP) for 16 h at 6 °C. The resulting solution was passed over Ni-NTA resin column to remove the un-cleaved protein and further purified by FPLC with HiLoad Superdex 200 (GE Biosciences) size exclusion column. Pure tag-free FabB was collected and concentrated to 2-4 mg/mL using Amicon Ultra Centrifuge Filters (MilliporeSigma) with 10 kDa molecular weight cutoff.

Native ACP recombinant protein was expressed in *E. coli* BL21(DE3) and cultured in LB broth containing 100 mg/L ampicillin. Cells were grown at 37 °C until OD_600_ reached ~0.7 and induced by 0.5 mM IPTG for 16 h at 18 °C. Cell pellets were collected by centrifugation, lysed by sonication in lysis buffer (50 mM Tris pH 7.4, 150 mM NaCl, 10% glycerol), and underwent another centrifugation at 17,400 g for 1 h to clear the lysate. Taking advantage of high ACP stability under 50% isopropanol solution, irrelevant proteins were precipitated by dripping in same volume of isopropanol into the lysate at the speed of 0.1 mL per second and removed by centrifugation at 17,400 g for 1 h. The supernatant was purified by FPLC via HiTrap Q HP anion exchange column with a linear gradient of 0 M to 1 M NaCl buffer. ACP eluted around 0.3 M NaCl and was collected to go through the second round of FPLC purification with HiLoad Superdex 75 PG (GE Biosciences) size exclusion column. Pure ACP was collected and concentrated using Amicon Ultra Centrifuge Filters (MilliporeSigma) with 3 kDa molecular weight cut off.

Actinorhodin KS (actKS) was prepared as described previously (Taguchi *et al*., 2017).

### 2.3. ACP modification and crosslinking reaction

ACP is overexpressed in a 1-to-1 ratio of the *apo* and *holo* form. Since the crosslinking probes contain the PPant moiety, the *apo*-ACP is the desired form, and thus the PPant on *holo*-ACP is removed by enzymatic reaction of the ACP hydrolase, AcpH. 2-10 mg/mL of ACP mixture can be totally converted to *apo*-ACP with 0.01 mg/mL of AcpH in the reaction buffer (50 mM Tris pH 7.4, 150 mM NaCl, 10% glycerol, 10 mM MgCl_2_, 5 mM MnCl_2_, 0.25% DTT) under 25 °C in 16 h. The resulting *apo*-ACP is purified by size exclusion (HiLoad Superdex 75 PG SEC) and concentrated (Amicon Ultra Centrifuge Filters, 3 kDa MWCO) to 2-10 mg/mL. To load the crosslinking probes onto *apo*-ACP, they were first converted to CoA analogues through the activities of three CoA biosynthetic enzymes (CoaA, CoaD, and CoaE) and transferred onto the serine residue (S36) of *apo*-ACP by the phosphopantetheinyl transferase Sfp. The whole modification and loading can be done in a one-pot chemoenzymatic reaction with the following condition: 1 mg/mL *apo*-ACP, 0.04 mg/mL Sfp, 0.01 mg/mL each CoA enzymes, and 0.2 mM crosslinking probe in reaction buffer (50 mM potassium phosphate pH 7.4, 12.5 mM MgCl_2_, 1 mM DTT) at 37 °C for 16 h. The stock solution of the probes was prepared by dissolving the probes in DMSO to a final concentration of 50 mM for C14Cl and 25 mM for C16:1Br due to the lower solubility of the longer acyl chain. The probe-loaded ACP, or *crypto*-ACP, was then purified by size exclusion and concentrated as above to 2-5 mg/mL. All the different forms of ACP can be tracked by the conformationally sensitive urea PAGE.

The crosslinking reaction was set up by mixing *crypto-ACP* with FabB in 3:1 ratio in reaction buffer (50 mM Tris pH 7.4, 150 mM NaCl) at 37 °C for 16 h. The reaction was monitored by 12% SDS PAGE and purified by size exclusion with minimal buffer (12.5 mM Tris pH 7.4, 25 mM NaCl). The crosslinked complex was concentrated to 8-10 mg/mL using Amicon Ultra Centrifuge Filters with 30 kDa MWCO and flash-frozen to store under −80 °C if not immediately used for crystallography.

### 2.4. Protein crystallography, data processing, and structure refinement

Crystals of all crosslinked complexes were grown by vapor diffusion in a cold room kept at 6 °C. 1 μL of crosslinked complex (8–10 mg·mL^-1^) was mixed with 1 μL of corresponding crystallographic condition and the mixture was placed inverted over 500 μL of the well solution (hanging-drop method). The FabB=C14Cl-ACP and FabB=C16:1-ACP crystals were grown in 18-24% (w/v) PEG 8 K, 0.2 M Mg(OAc)_2_, 0.1 M sodium cacodylate pH 6.5. These conditions produced square plates and required two weeks for complete crystal growth. X-ray diffraction data (Table S1) were collected at the Advanced Light Source synchrotron at Berkeley. Data were indexed using iMosflm (Battye *et al*., 2011) then processed and scaled using aimless from the CCP4 software suite (Winn *et al*, 2011). Scaled reflection output data were used for molecular replacement and model building in PHENIX (Adams *et al*., 2010). Initial phasing via molecular replacement was performed by searching for the full FabB=ACP complex using 5KOF as a search model. Subsequent refinement and model refinement in PHENIX and Coot, respectively. The parameter files for the covalently bonded 4’-phosphopantetheine were generated using Jligand (Lebedev *et al*, 2012). Manually programmed parameter restraints were used to create the associated covalent bonds between 4’-phosphopantetheine to Ser36 of ACP and Cys163 of FabB during refinement.

## 3. Results and discussion

### 3.1. Rationale behind crosslinker choice and the KS stereospecificity for chloroacrylate probes

To understand the molecular basis of FabB substrate preference toward long chain FAs, we developed a 14-carbon chloroacrylate crosslinker, C14Cl, that we hypothesized would mimic the condensation step between the longest favoured FabB substrate, C12, and malonyl-ACP. Probes containing two different configurations, *E* and *Z*-form, respectively, of the acrylate double bond were synthesized and tested (Fig. S1). Only the *Z*-form showed crosslinking activity on FabB (Fig. S2A) and the same applies to the other two tested KSs, FabF and actinorhodin KS (actKS), a type II polyketide synthase (Keatinge-Clay *et al*., 2004). Given that there are published crosslinked structures of FabB-ACP and FabF-ACP utilizing (*E*)-C3Cl and (*E*)-C8Cl, respectively, we hypothesized that while *Z*-form is the preferred configuration, the selectivity only develops at longer chain lengths due to the restrictions in the substrate pocket. To test this hypothesis, we performed crosslinking assays using (*E*)-C8Cl and actKS, which has a narrower pocket more likely to clash with the acyl substrate. Results from these assays showed no crosslinking activity (Fig. S2B). However, (*E*)-C3Cl was able to crosslink with our entire panel of KSs, indicating that actKS is generally active towards unsubstituted *E*-form chloroacrylate crosslinkers. As a result, the data suggest stereospecificity of KSs toward the *Z*-form long-chain chloroacrylate crosslinker. After confirming the preferred stereochemistry, we loaded ACP with (*Z*)-C14Cl and performed large-scale crosslinking to obtain the FabB=C14-ACP for structural analysis.

To explore mechanisms governing FabB substrate selectivity, we also obtained a crosslinked complex using the C16:1-α-bromo crosslinker, or C16:1Br (Fig. S1), likely to mimic the transacylation step of an unfavoured substrate, namely C16:1-ACP. Importantly, this substrate is readily accepted by FabF *in vitro* and *in vivo*, and its elongation to C18:1 is critical for *E. coli’s* rapid homeoviscous adaptive response to regulate membrane fluidity due to changes in environmental temperature (Garwin *et al*., 1980; de Mendoza & Cronan, 1983). Previously, we utilized C16:1-ACP to investigate the molecular mechanisms governing FabF’s unique capacity to extend C16:1 to C18:1 (Mindrebo *et al*., 2021). A FabB=ACP structure using the same crosslinker could potentially inform how FabB discriminates against longer acyl substrates.

### 3.2. FabB-ACP trapped in different catalytic states by the crosslinkers

The FabB=C14-ACP complex was crystallized and diffracted to a nominal resolution of 1.7 Å. The asymmetric unit contains the biologically relevant dimeric structure (FabB-ACP)_2_ with well-defined electron density for the crosslinker (Fig. 1). The active site cysteine, C163, is covalently attached to the C3 position of the acyl chain, which resembles the number of carbons of the putative tetrahedral intermediate formed during the condensation half-reaction. The carbonyl group of the acyl chain coordinates with the two active site catalytic histidine residues, H298 and H333, as seen in all other KS-ACP crosslink structures using the chloroacrylate crosslinkers (Milligan *et al*., 2019; Du *et al*., 2020; Mindrebo *et al*., 2021). Interestingly, the double bond of the acrylate is in the same *trans* configuration as observed in other structures despite the use of *E*-form crosslinkers. This suggests a stereoselective addition-elimination reaction mechanism governed by the active site arrangement during reformation of the double bond. Overall, the two protomers of the structure align with an RMSD of 0.872 Å and show no obvious structural differences.

**Figure 1.**
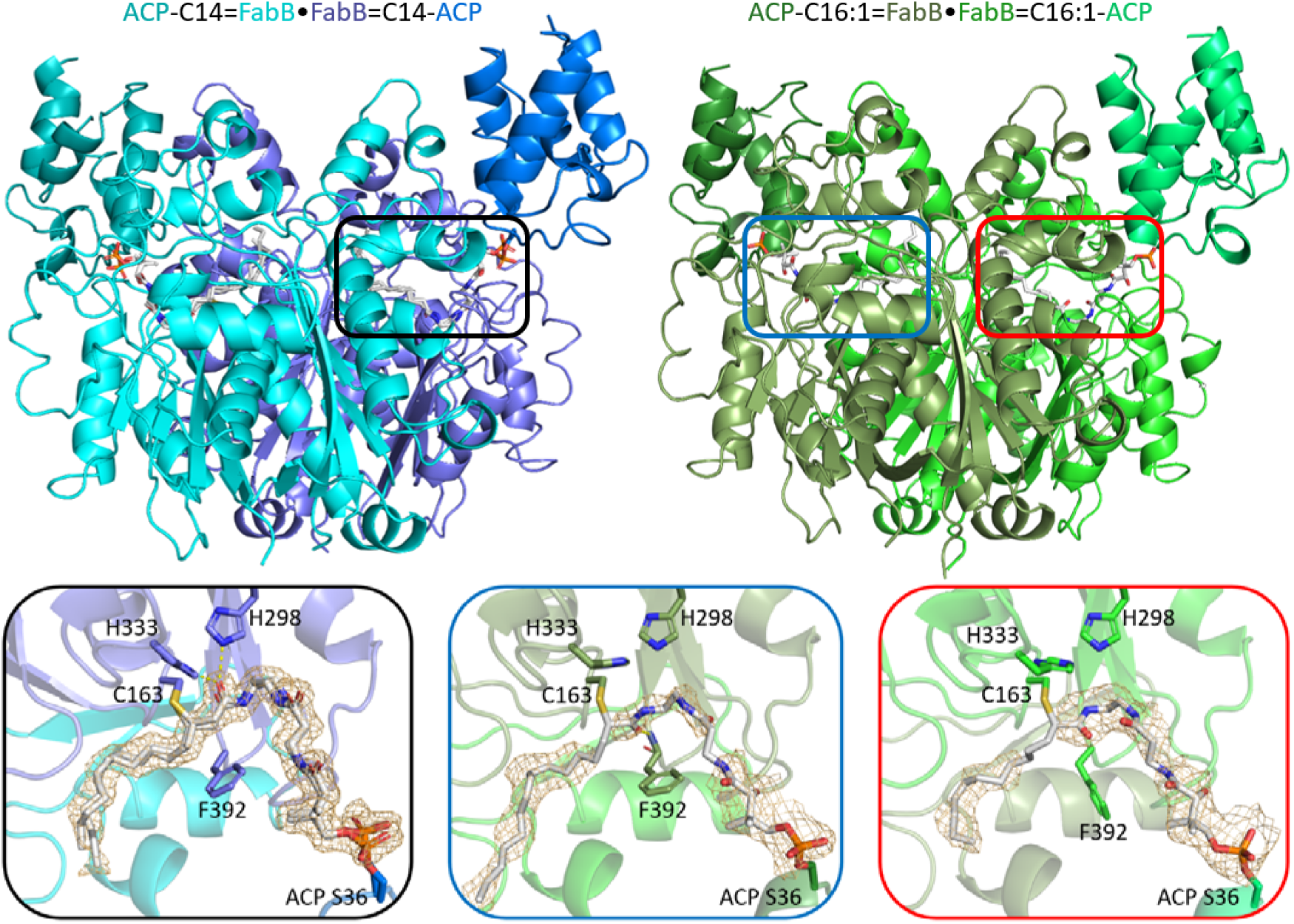
FabB trapped in different states by crosslinking with ACP. FabB=C14-ACP (PDB ID: 7SQI), blue structure on the left, possesses two identical active sites with the crosslinker solved in alternative conformations. One of its active sites is enlarged in the black frame, which depicts a condensation half-reaction arrangement. FabB=C16:1-ACP structure (green, PDB ID: 7SZ9) on the right has two distinct protomers. Protomer A (blue frame) depicts a transacylation half-reaction arrangement while protomer B (red frame) features partially solved aliphatic chain and a catalytic incompetent F392 rotamer, potentially mimicking a substrate release state. The mesh surface depicts the Fo-Fc polder omit map calculated by omitting the crosslinker (contoured at sigma value 3.0, 2.0, and 3.0, respectively, from left to right).

The FabB=C16:1-ACP complex was crystallized and diffracted to 2.20-Å resolution, and the asymmetric unit contains the same dimeric arrangement (Fig. 1). The protomer that contains the chain A monomer of FabB, referred to as protomer A, has the acyl chain carbonyl positioned in the oxyanion hole formed by the backbone amides of C163 and F392. This coordination is essential for the transacylation half-reaction (Olsen *et al*., 2001; Rittner *et al*., 2020). The other protomer of the complex, referred to as protomer B, captures a unique active site arrangement where the acyl carbonyl does not form polar contacts with any residue, but positions itself so that it repels F392 away from the active site, disassembling the oxyanion hole. Importantly, F392 is a crucial gating element and its rotamers regulate malonyl substrate entry (Zhang *et al*., 2006; Luckner *et al*., 2009; Witkowski *et al*., 1999). Furthermore, it also plays a critical role in the KS gating mechanism during substrate delivery and undergoes an ~8 Å residue shift (Cα to Cα) when transitioning to the gate-open conformation. Due to the unfavourable substrate conformation, protomer B resembles a catalytically unproductive state (oxyanion hole not present) with an altered F392 gating residue position. It is possible that this conformation represents an active site primed for substrate release. An overlay of the two protomers with our recently published gate-open FabF=C16:1-ACP (PDB ID: 7L4E) structure suggests a potential trajectory for gate opening, starting from the gate-closed protomer A to the disrupted protomer B and finally the gate-open FabF=ACP structure (Fig. 2). The proposed transition features a 90° rotation of the F392 side chain, likely driven by the acyl carbonyl flipping out from the oxyanion hole, which places the side chain underneath the PPant arm to facilitate the subsequent transition. The loop then undergoes a large conformational change to the gate-open structure, with G391 and G394 acting as hinges, creating space for product dissociation. During the gate transition, loop 2 undergoes a conformational change in concert with loop 1, as observed in FabF. The adjacent loop 2 residue, V270, abuts F392 and would sterically impede a transition to the gate open conformation. V270 from protomer B has a notably higher B-factor (135.5) than that of protomer A’s V270 (93.1), indicating a less ordered loop 2 potentially primed for transition to the open state. To better illustrate the gate transition, we used PyMOL to generate an interpolated trajectory between the three states in Figure 2 (Movie S1).

**Figure 2.**
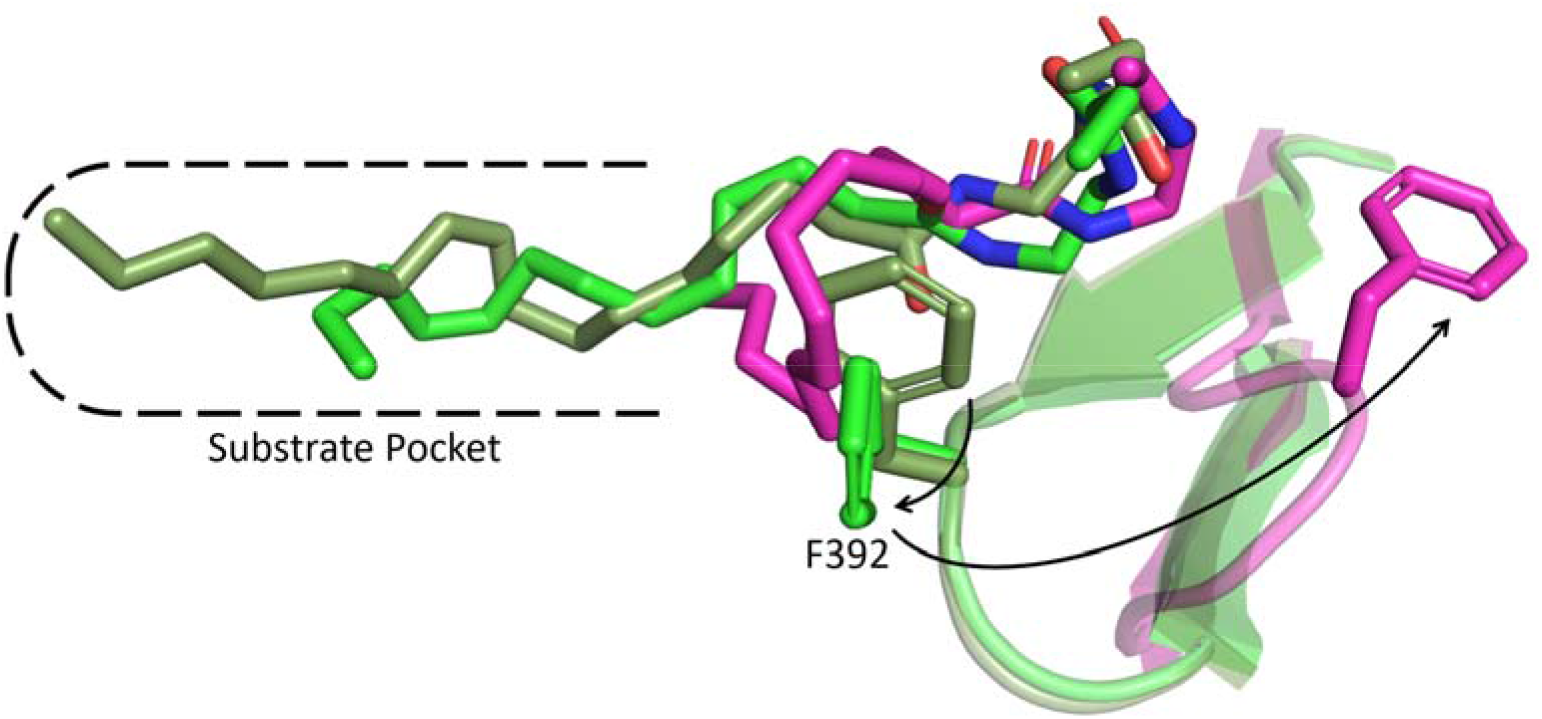
A potential trajectory for substrate de-sequestration is illustrated by overlaying three active sites from two crosslinked KS-ACP structures utilizing the same C16:1Br crosslinker. The arrow indicates the transition of the gating residue phenylalanine from protomer A (dark green, gate closed) to protomer B (light green), and finally the gate open form (magenta, FabF-ACP structure, PDB ID: 7L4E). An animation (Movie S1) with interpolated trajectories between the three states can be found in the supporting information.

### 3.3. Asymmetric pockets of the FabB homodimer suggest negative cooperativity

The substrate pocket of FabB extends from the active site cysteine C163 to the homodimer interface. At the interface, E200 and Q113’ (The apostrophe denotes the residue from the dyad-related protomer) sit near the bottom of the pocket forming side chain polar contacts that modulate the size of the binding pocket. However, these residues can also adopt different rotamers where the side chains turn away from each other in opposite directions, breaking the interaction and resulting in pocket expansion. This double side chain rotation can be classified as a “swinging door enzyme gate” (Gora *et al*., 2013), and will hereafter be referred to as the “E-Q gate”. In the FabB=C14-ACP structure, where a 12-carbon aliphatic chain extends into the binding pocket, two sets of alternative conformations were refined for both the acyl chains and E-Q gates. The E-Q gates from both pockets can be modelled in the open (expanded pocket) or closed (normal pocket) states. The open E-Q gate conformation places the Q113’ residue away from the pocket towards the dimer interface and expands the size of the binding pocket when compared to the closed state. However, the modelled C12 substrate conformations are readily accommodated by both E-Q gate conformations (Fig. 3A). Interestingly, there is only enough space at the dimer interface to accommodate Q113 from a single protomer, indicating that the two open E-Q gate conformations cannot coexist.

**Figure 3.**
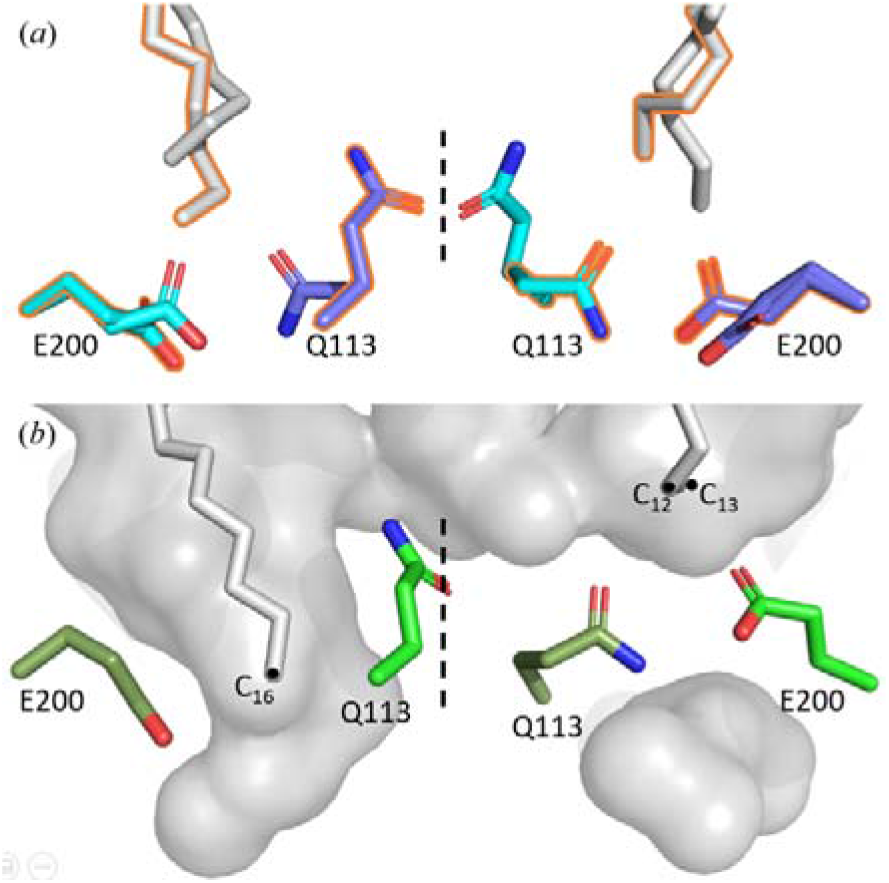
Asymmetric pockets and the E-Q gate. Residues with the same colour belong to the same FabB monomer. (*a*) FabB=C14-ACP possesses two sets (orange frame and no frame) of alternative conformations of the acyl chain and the E-Q gate. (*b*) FabB=C16:1-ACP has an expanded pocket (left, protomer A, open E-Q gate) and a normal pocket (right, protomer B, closed E-Q gate).

FabB’s asymmetric pocket features are further illustrated in the FabB=C16:1-ACP structure, in which protomer A has an expanded pocket and protomer B does not (Fig. 3B). In protomer A, the 16-carbon aliphatic chain penetrates through the open E-Q gate, with the last four carbons flanked by the side chains of the two gating residues. The expanded pocket extends to the α-helix (residue 110’-121’) that contains Q113’. Protomer B, on the other hand, has a closed E-Q gate that distorts the C16 chain, which prohibited the modelling of more than 13 carbons into the available density. As a result, a FabB binding pocket in the E-Q gate-closed conformation cannot comfortably accommodate an acyl chain longer than 12 carbons. Importantly, the sterically occluded chain in protomer B likely contributes to the distorted active site arrangement noted above. These observations suggest the inability to accommodate the acyl substrate results in improper positioning of the thioester moiety for catalysis, prohibiting the transfer of unfavourable substrates to the KS cysteine residue during the transacylation half-reaction.

FabB shows activity elongating FAs up to 14 carbons in length (Edwards *et al*., 1997). This is supported by the FabB=C14-ACP structural data, as both protomers can accommodate 12 carbon substrates regardless of the E-Q gate conformation. However, only the expanded pocket can readily accommodate acyl substrates beyond 12 carbons in length. These results explain the low, but existing, FabB activity towards long-chain acyl substrates *in vitro* and *in vivo* (Edwards *et al*., 1997; Mindrebo *et al*., 2020b). Furthermore, since only one E-Q gate can occupy the open conformation, these data suggest a negative cooperativity mechanism toward long-chain acyl substrates, as observed in the FabB=C16:1-ACP structure, where protomer B “struggles” to accept the substrate. Therefore, the observed pocket and E-Q gate mechanism align well with the reported activity of FabB.

### 3.4. The implication of the side pocket in KS mechanism

In FAS KSs, there is a “side pocket’ located adjacent to the catalytic active site and opposite the side of the substrate binding pocket. The PPant binding tunnel, acyl substrate binding cavity, and the side pocket form a “τ” shaped space (Fig. S3). The side pocket has a surface area of 180 Å^2^ and a volume of 95 Å^3^, which is larger than the acyl substrate pocket that has a surface area of 145 Å^2^ and a volume of 58 Å^3^, as calculated by CASTp (Tian *et al*., 2018). Despite its proximity to the active site, the side pocket has only been examined in passing in previous studies and has not been assigned a role in catalysis. The recently discovered gating loops reveal that the side pocket, which is partially formed by loop 1 and loop 2 when in the gate-closed conformation, is required to accommodate the loop 1 phenylalanine gating residue in the gate-open conformation (Fig. S4). Introducing bulky residues by site-directed mutagenesis that fill in the available side-pocket space significantly impairs FabF activity (Mindrebo *et al*., 2021) by restricting access to the gate-open conformation.

Interestingly, the high resolution FabB=C14-ACP structure allows us to unambiguously assign 10 water molecules in the side pocket that form a water-mediated network connecting loop 1, loop 2, and the PPant moiety (Fig. 4). In the proximity of the catalytic triad, two water molecules, WAT1 and WAT2 (Fig. S5), abut the catalytic triad, forming another water-mediated network linking T300, D306, and E309. Notably, these three residues are absolutely conserved across 461 FabB sequences spanning bacterial species. Previous work suggests that a nucleophilic water molecule, activated by the catalytic histidine residue (H298 in FabB), is required for malonyl-ACP decarboxylation in order to form the enolate anion that subsequently attacks the acyl-KS intermediate. In this mechanism, a hydroxide ion attacks the malonyl carboxylate moiety to form bicarbonate as the released product instead of CO_2_ (Zhang *et al*., 2006; Witkowski *et al*., 2002). The observed water network in the side pocket creates a hydrophilic and water-rich site adjacent to the reaction chamber. This network might assist the decarboxylation of malonyl-ACP by providing such a water molecule needed for bicarbonate formation, as well as for accommodating the bicarbonate product. This observation supports the bicarbonate-forming mechanism, as opposed to the mechanism that posits release of carbon dioxide (Chisuga *et al*., 2022). More biochemical and structural studies are required to definitively address this long-standing question (Heil *et al*., 2019).

**Figure 4.**
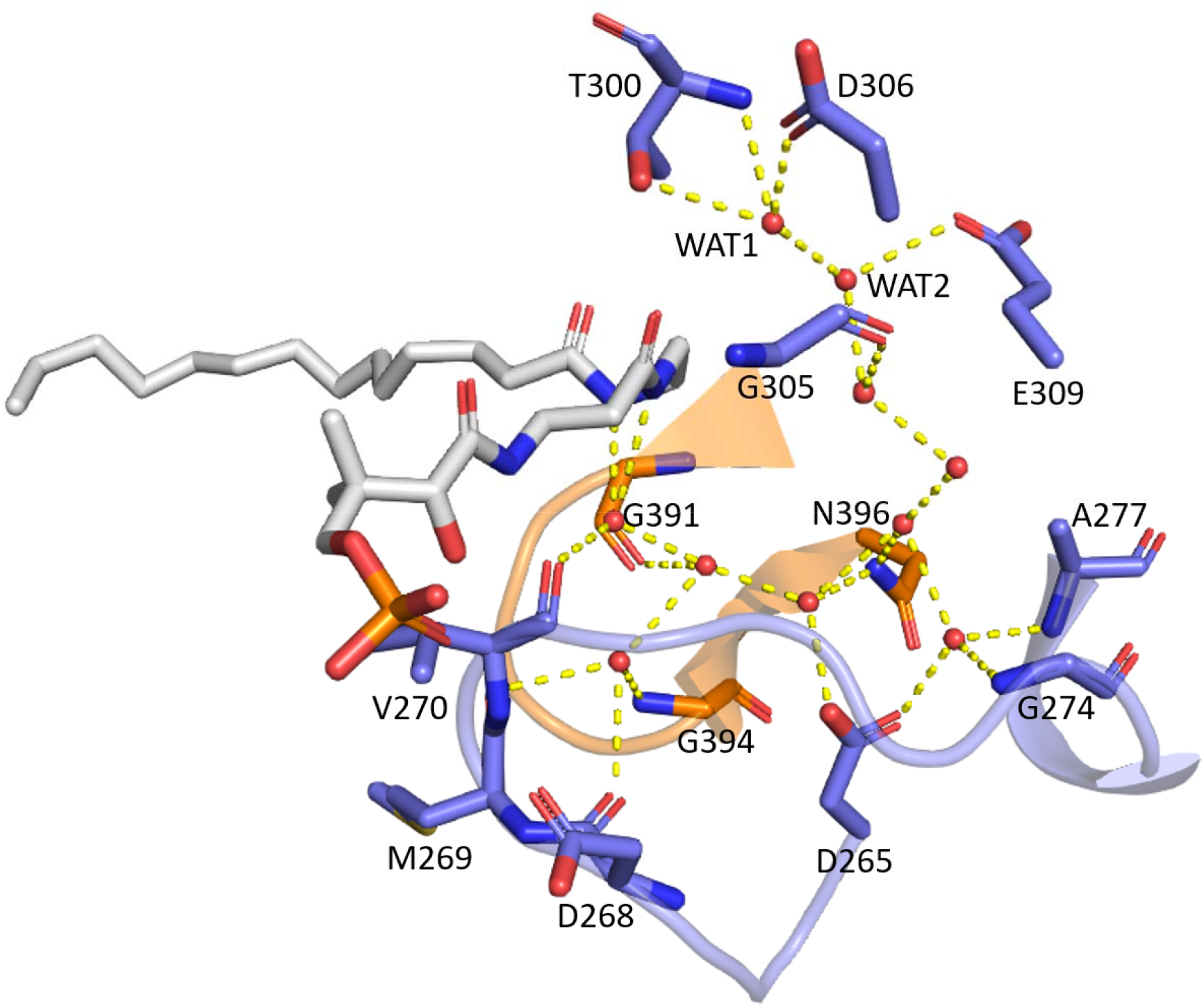
Ten water molecules in the side pocket form a water network between highly conserved residues and the PPant arm. Cartoon representation is shown for the gating loop 1 (orange) and loop 2 (blue). All polar contacts (yellow dashed line) are within 3.2 Å.

## 4. Conclusion

Since its initial discovery (Cronan *et al*., 1969), *E. coli* FabB has been studied for more than 50 years, with at least 17 structures published and deposited with the Protein Data Bank (*apo*, mutants, substrate-bound, inhibitor-bound, and ACP-crosslinked). Despite the extensive knowledge we have regarding this enzyme, some fundamental questions, such as substrate specificity and the exact decarboxylation mechanism, have yet to be fully understood. In this study, we leveraged our understanding of the KS reaction mechanism and applied mechanism-based crosslinkers to elucidate two FabB=ACP crosslinked complexes in both favoured and disfavoured catalytic states. The C14Cl crosslinker captures FabB in an on-pathway state, representing the condensation reaction with a favoured substrate, C12, and malonyl-ACP. Alternatively, the complex trapped using the C16:1Br crosslinker represents the acylation reaction with a disfavoured long-chain acyl-ACP substrate. Analysis of these structures led to the identification of an E-Q gate at the back of the substrate binding pocket that enforces asymmetry between the two protomers. A 12-carbon substrate can be readily accommodated in both pockets. However, the E-Q gate must transition to the open conformation to accommodate longer fatty acid substrates to appropriately position the thioester moiety for the transacylation reaction. The asymmetric nature of the E-Q gate suggests negative cooperativity of FabB towards long chain substrates, which may explain the reduced activity of FabB towards C14 and C16:1, two substrates readily extended by FabF. In addition to these mechanistic insights, we have characterized a water-rich side pocket that plays an important role in the dual-loop KS gating mechanism and supports the bicarbonate-forming decarboxylation mechanism. Results from these studies provide important insights into *E. coli* FAS and broadly inform mechanisms governing KS-mediated catalysis.

**Table 1.** X-ray crystallography data collection and refinement statistics.

| Structure (PDB ID) | FabB=C14-ACP (7SQI) | FabB=C16:1-ACP (7SZ9) |
| --- | --- | --- |
| Wavelength (Å) | 1 | 1 |
| Resolution range (Å) | 36.86 - 1.7 (1.761 - 1.7) | 47.9 - 2.2 (2.279 - 2.2) |
| Space group | P 21 21 21 | P 1 21 1 |
| Unit cell (Å, °) | 58.99 112.38 141.64 90 90 90 | 59.4505 103.58 79.2802 90 108.28 90 |
| Unique reflections | 104222 (10240) | 38076 (2494) |
| Reflections used in refinement | 103901 (10155) | 37907 (2488) |
| Reflections used for R-free | 5103 (466) | 1808 (122) |
| R-work | 0.1608 (0.3039) | 0.2084 (0.3117) |
| R-free | 0.1981 (0.3281) | 0.2553 (0.3461) |
| Protein residues | 965 | 957 |
| Solvent | 1069 | 165 |
| RMS (bonds) | 0.008 | 0.002 |
| RMS (angles) | 0.97 | 0.47 |
| Ramachandran favoured (%) | 96.33 | 94.84 |
| Ramachandran allowed (%) | 3.36 | 4.43 |
| Ramachandran outliers (%) | 0.31 | 0.74 |
| Rotamer outliers (%) | 0.80 | 1.46 |
| Clashscore | 3.80 | 4.84 |
| Average B-factor | 23.68 (overall)<br>24.48 (protein)<br>19.97 (crosslinker)<br>30.89 (water) | 57.06 (overall)<br>57.11 (protein)<br>78.61 (crosslinker)<br>45.80 (water) |

## Acknowledgements

This work was supported by NIH R01 GM095970. J. T. M. was supported by T32 GM832626. T. D. D. was supported by NIH K12 GM068524 and K99 GM12945.

## Supporting information

### A. Supporting figures

**Figure S1.**
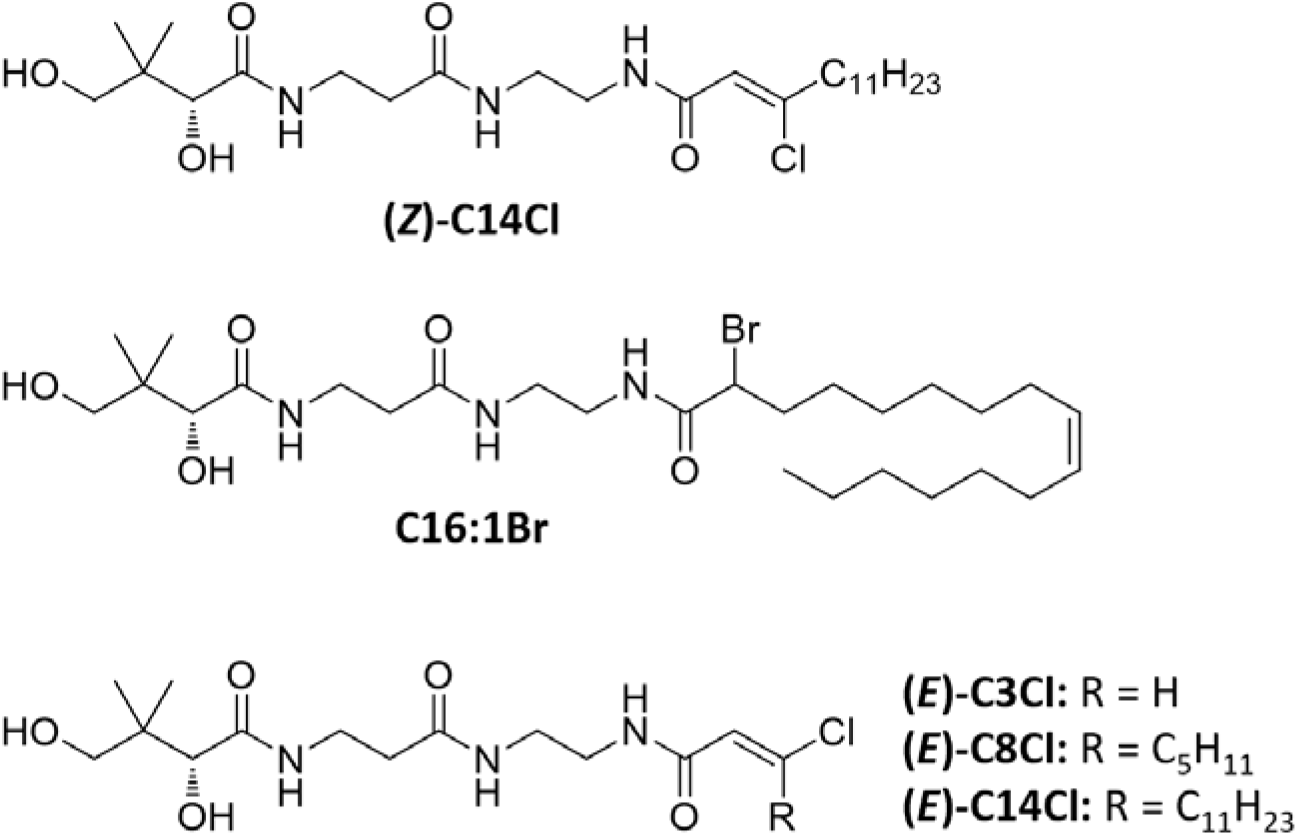
Crosslinkers used in this study. The top two crosslinkers were used to obtain the two FabB-ACP crosslinked structures while the tree *E* form crosslinkers were used for crosslinking activity study.

**Figure S2.**
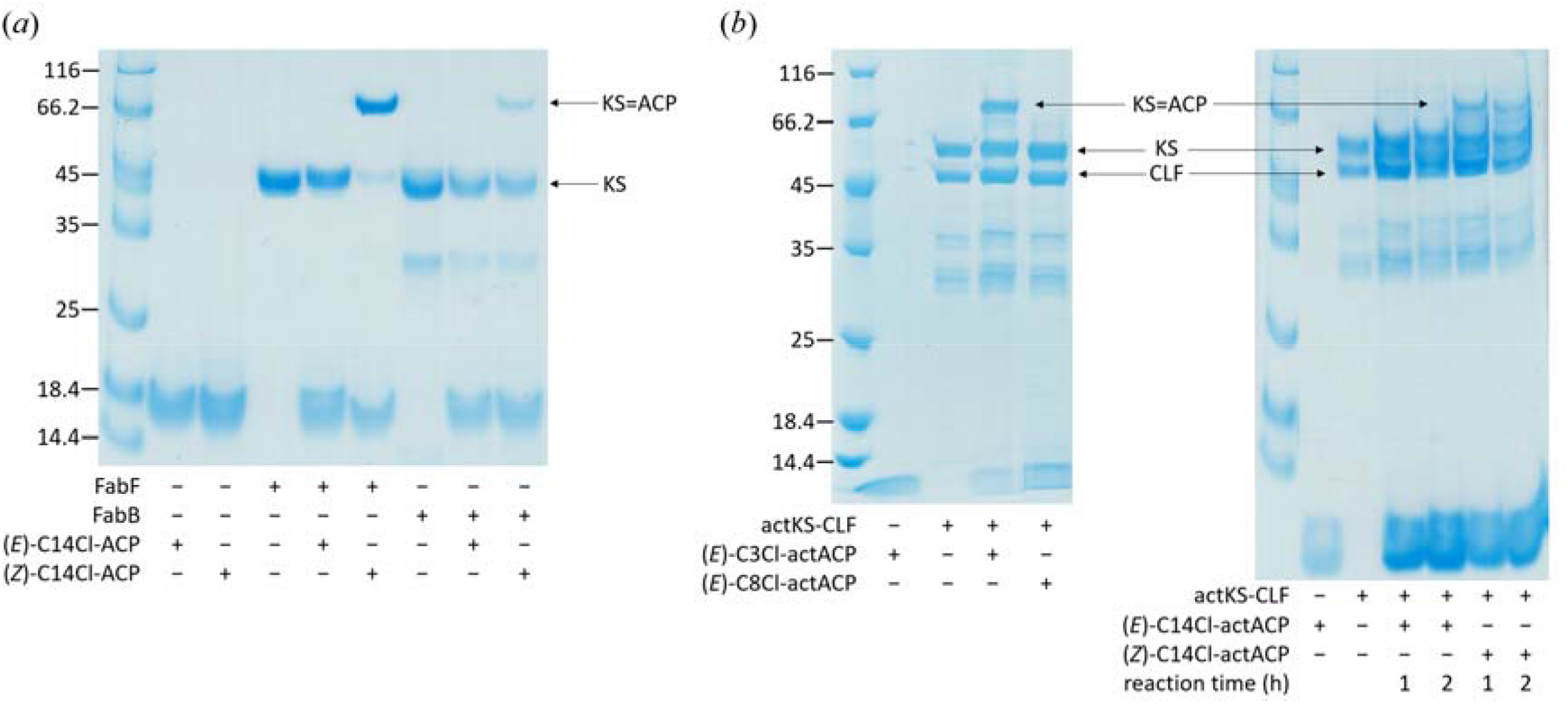
Crosslinking test on three different KSs shows stereospecificity toward *Z*-form chloroacrylate crosslinker. Samples were run on 12% SDS-PAGE and stained by Coomassie Brilliant Blue. (*a*) (*Z*)-C14Cl can crosslink ACP with FabF or FabB whereas (*E*)-C14Cl shows no crosslinking activity. (*b*) Actinorhodin ketosynthase (actKS-CLF) exists as a heterodimer, in which the chain length factor subunit lacks the active site residues and is slightly smaller than KS subunit in size. Only (*E*)-C3Cl and (*Z*)-C14Cl show crosslinking activity toward actKS-CLF and actACP.

**Figure S3.**
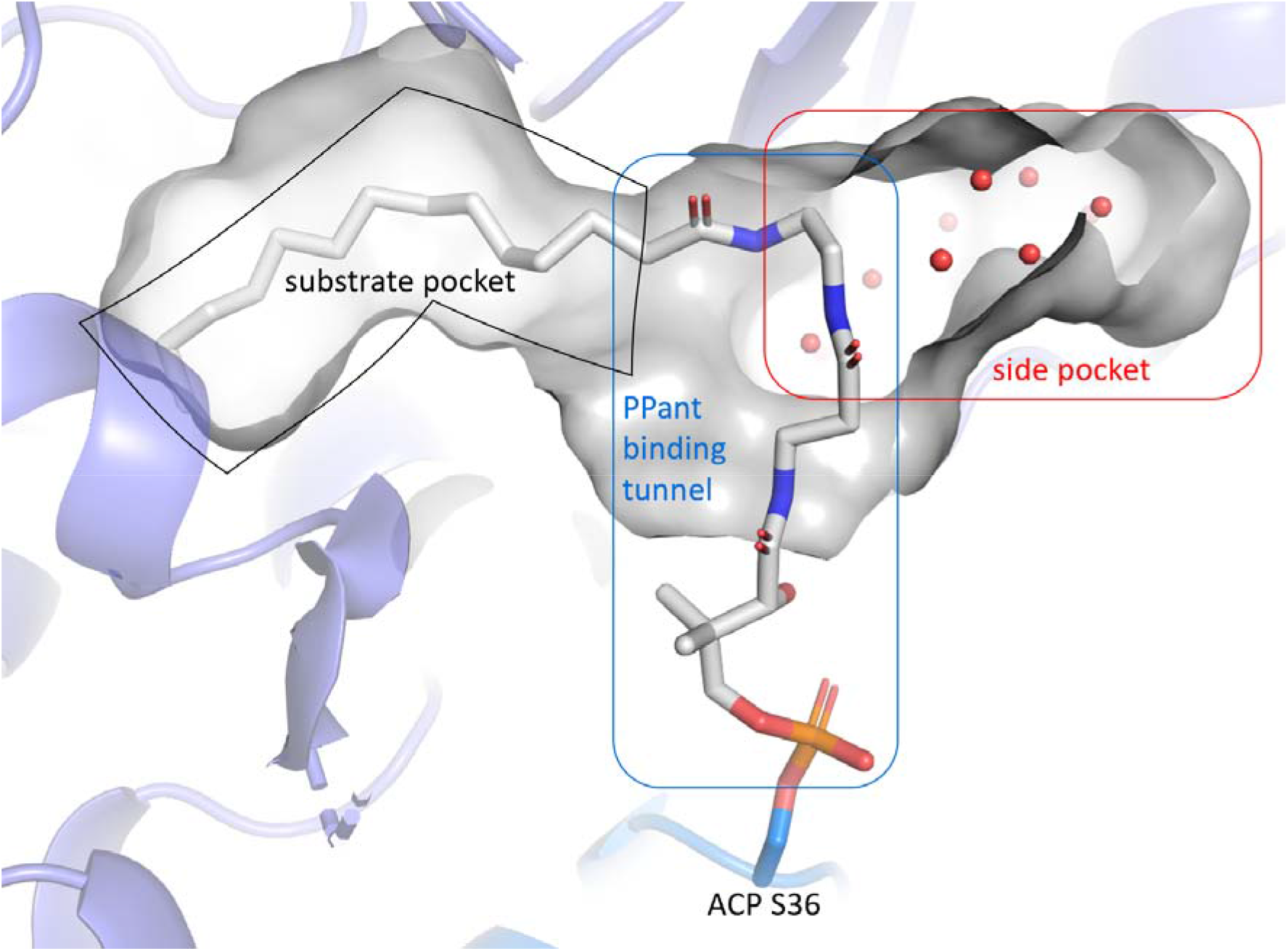
PPant binding tunnel, substrate pocket, and the side pocket form a “τ” shaped space inside FabB. This figure is generated from the FabB=C14-ACP (PDB ID: 7SQI) structure.

**Figure S4.**
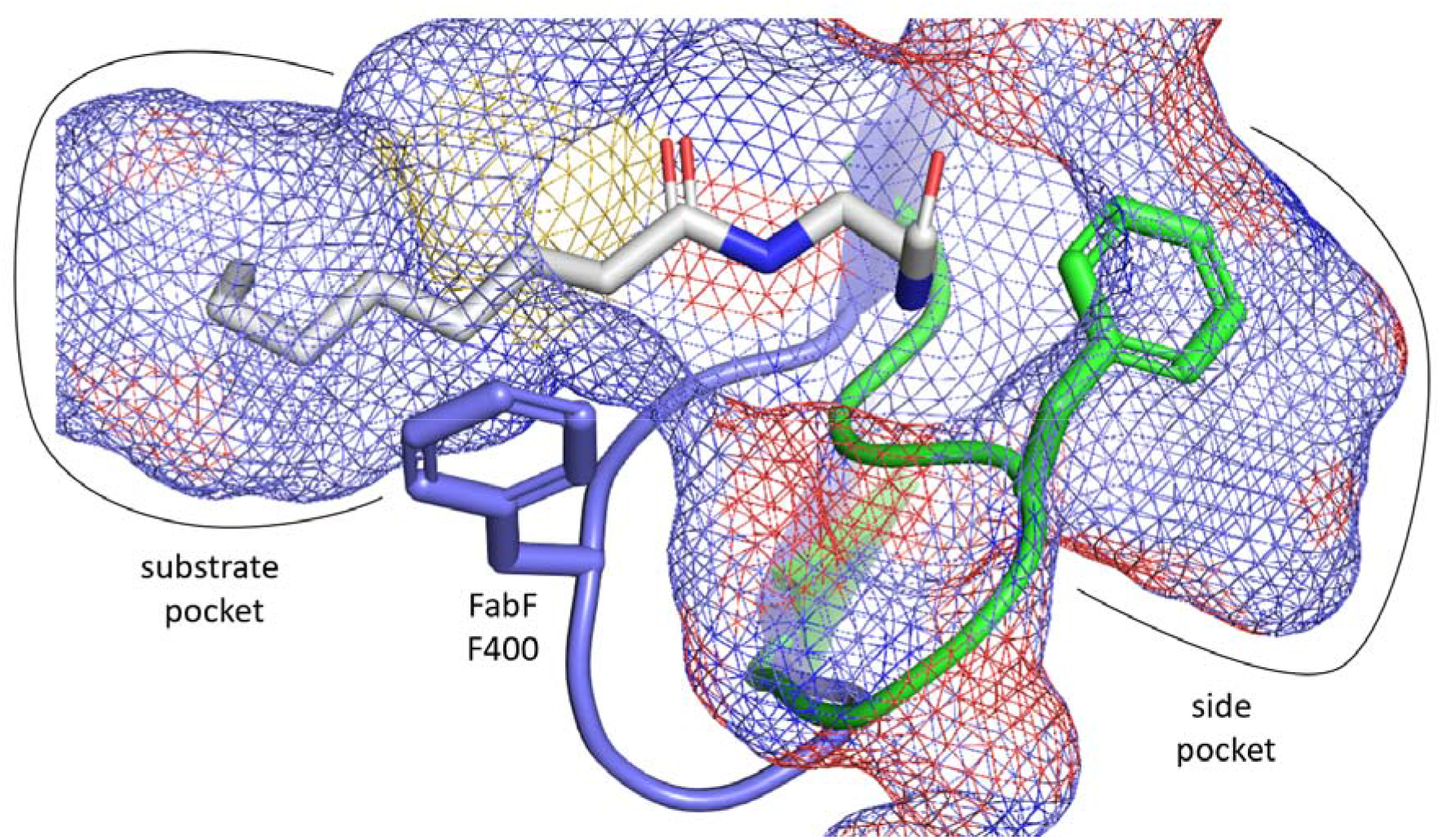
Overlay of the open form gating loop 1 (green, PDB ID: 6OKG) with the structure in closed form (blue, PDB ID: 7L4L). The side pocket accommodates the open loop.

**Figure S5.**
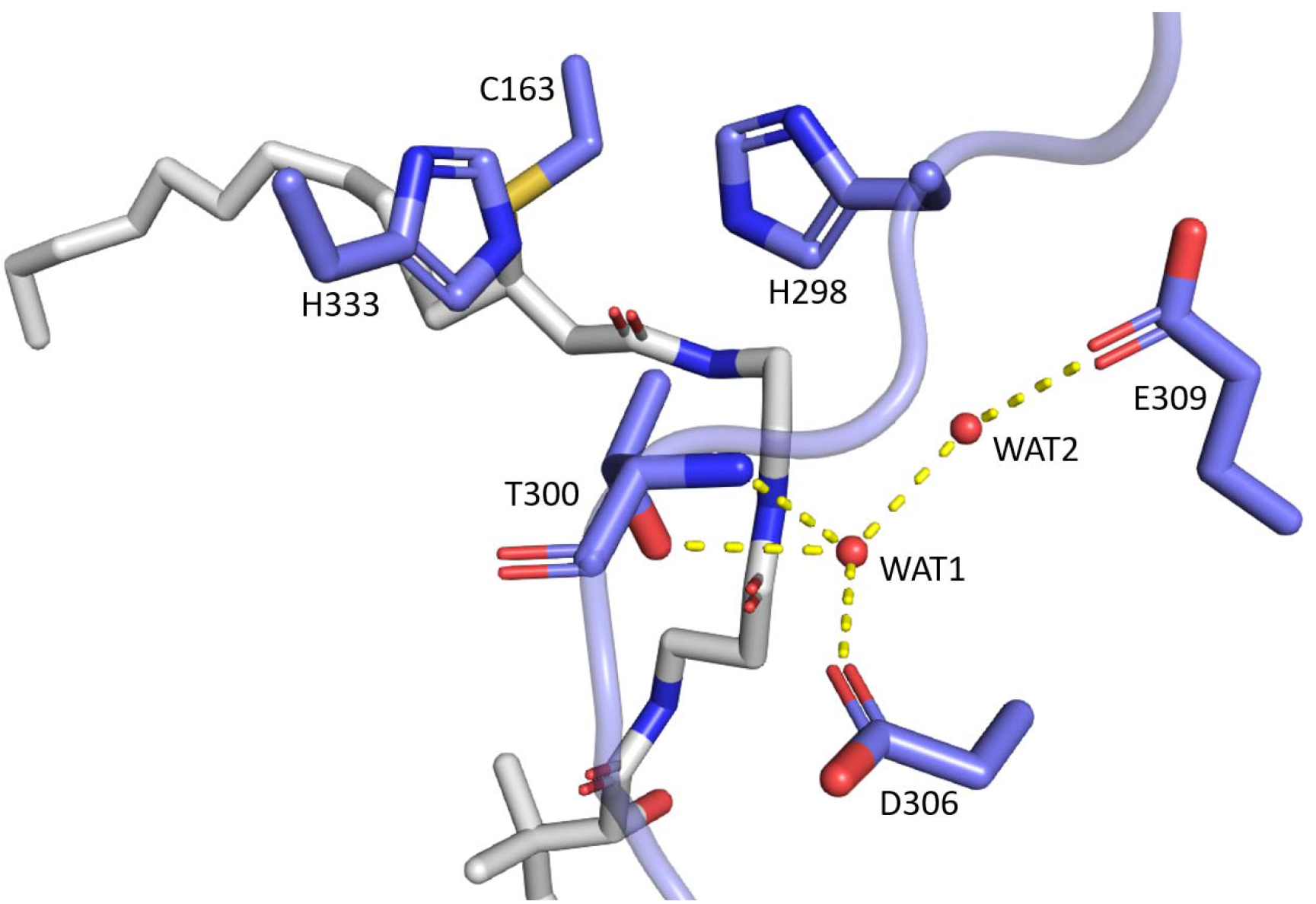
The two water molecules, WAT1 and WAT2, respectively, in proximity of the catalytic histidine residues. All residues shown are 100 % conserved over 461 FabB sequences aligned.

### B. Synthesis of crosslinking probes

Chemical reagents were purchased from Acros, Fluka, Sigma-Aldrich, or TCI. Deuterated NMR solvents were purchased from Cambridge Isotope Laboratories. All reactions were conducted with vigorously dried anhydrous solvents that were obtained by passing through a solvent column exposed of activated A2 alumina. All reactions were performed under positive pressure of argon in flame-dried glassware sealed with septa and stirred with Teflon coated stir bars using an IKAMAG TCT-basic mechanical stirrer (IKA GmbH). Analytical Thin Layer Chromatograpy (TLC) was performed on Silica Gel 60 F254 precoated glass plates (EM Sciences). Visualization was achieved with UV light and/or appropriate stain (I_2_ on SiO_2_, KMnO_4_, bromocresol green, dinitrophenylhydrazine, ninhydrin, or ceric ammonium molybdate). Flash column chromatography was carried out with Geduran Silica Gel 60 (40–63 mesh) from EM Biosciences. Yield and characterization data correspond to isolated, chromatographically, and spectroscopically homogeneous materials. ^1^H NMR spectra were recorded on Varian Mercury 400, Varian Mercury Plus 400, or JEOL ECA500 spectrometers. ^13^C NMR spectra were recorded at 100 MHz on Varian Mercury 400 or Varian Mercury Plus 400 spectrometers. Chemical shifts for ^1^H NMR and ^13^C NMR analyses were referenced to the reported values (Gottlieb *et al*., 1997) using the signal from the residual solvent for ^1^H spectra, or to the ^13^C signal from the deuterated solvent. Chemical shift δ values for the ^1^H and ^13^C spectra are reported in parts per millions (ppm) relative to these referenced values, and multiplicities are abbreviated as s=singlet, d=doublet, t=triplet, q-quartet, m=multiplet, b=broad. All ^13^C NMR spectra were recorded with complete proton decoupling. FID files were processed using MestreNova 10.0 (MestreLab Research). Electrospray ionization (ESI) mass spectrometric analyses were preformed using a ThermoFinnigan LCQ Deca spectrometer. Spectral data and procedures are provided for all new compounds and copies of spectra have been provided.

#### 2-tetradecynoic acid (S1)

In a 50 mL round-bottomed flask, 1-tridecyne (0.12 mL, 0.52 mmol, 1.0 equiv.) and 5 mL THF were added. The vessel was cooled to 0 °C before the slow, dropwise addition of nBuLi (1.6 M in hexanes, 0.40 mL, 0.64 mmol, 1.1 equiv.). The reaction was stirred for 2 hours at 0 °C before flushing with carbon dioxide gas. After stirring for 3 hours at 0 °C, the reaction was warmed to room temperature (20 °C) and stirred overnight for 17 hours. The reaction was quenched with saturated aqueous NH_4_Cl (10 mL) and extracted with ethyl acetate (4×25 mL). The combined organic extracts were dried (MgSO_4_), filtered, and concentrated by rotary evaporation. Purification by silica flash chromatography (9:1 hexanes/ethyl acetate → 3:2 hexanes/ethyl acetate + 1% acetic acid) afforded carboxylic acid **S1** (96.7 mg, 83%, white solid).

**TLC:** R_f_ 0.21 (4:1 hexanes/ethyl acetate + 1% acetic acid). **^1^H-NMR** (400 MHz, CDCl_3_): δ10.98 (s, 1H), 2.34 (t, *J* = 7.0 Hz, 1H), 1.81–1.55 (m, 2H), 1.39 (s, 3H), 1.25 (s, 14H), 0.87 (t, *J* = 5.6 Hz, 3H).**^13^C-NMR** (100 MHz, CDCl_3_): δ 158.83, 92.92, 72.77, 32.03, 29.71, 29.53, 29.13, 28.94, 27.51, 22.81, 19.01, 18.88, 18.74, 14.19.

#### 3-chloro-2-tetradecenoic acid (S2)

In a 10 mL pear-shaped flask, 2-tetradecynoic acid **S1** (126.1 mg, 0.5621 mmol, 1.0 equiv.) and 5 mL DMF were added. The vessel was cooled to 0 °C before the addition of SOCl_2_ (48.9 μL, 0.675 mmol, 1.2 equiv.). After stirring for 1 hour at 0 °C, the reaction was warmed to room temperature (20 °C). After 3 hours at room temperature, additional SOCl_2_ was added (20 μL) and stirring was continued overnight. After 15 hours, additional SOCl_2_ was added (48.9 μL). After 2 hours, the reaction was concentrated by rotary evaporation, and the resulting liquid was poured over 12 grams of ice and extracted with diethyl ether (4×25 mL). The combined organic extracts were dried (MgSO_4_), filtered, and concentrated by rotary evaporation. Purification by silica flash chromatography (9:1 hexanes/diethyl ether → 4:1 hexanes/diethyl ether) afforded (*E*)-3-chloro-2-tetradecenoic acid **S2a** (39.2 mg, yellow solid) and (*E*)-3-chloro-2-tetradecenoic acid **S2a** (57.2 mg, brown solid)

#### (*E*)-3-chloro-2-tetradecenoic acid (S2a)

**TLC:** R_f_ 0.61 (1:1 hexanes/diethyl ether). **^1^H-NMR** (400 MHz, CDCl_3_): δ 6.09 (s, 1H), 2.98 (t, 2H), 1.63 (m, 2H), 1.26 (s, 14H), 0.95 – 0.80 (t, 3H).**^13^C-NMR** (100 MHz, CDCl_3_): δ 170.11, 161.06, 118.44, 36.11, 32.08, 29.78, 29.62, 29.51, 29.45, 28.90, 27.83, 22.86, 14.27.

#### (*Z*)-3-chloro-2-tetradecenoic acid (S2b)

**TLC:** R_f_ 0.45 (3:2 hexanes/diethyl ether). **^1^H-NMR** (400 MHz, CDCl_3_): δ 6.05 (s, 1H), 2.46 (t, *J* = 7.4 Hz, 1H), 1.63 (m, 2H), 1.26 (s, 15H), 0.88 (t, 3H).**^13^C-NMR** (100 MHz, CDCl_3_): δ 169.52, 154.24, 115.60, 41.74, 32.05, 29.73, 29.58, 29.47, 29.38, 28.67, 27.35, 22.83, 14.25.

#### (4*R*)-*N*-(3-((2-((*E*)-3-chlorotetradec-2-enamido)ethyl)amino)-3-oxopropyl)-2-(4-methoxyphenyl)-5,5-dimethyl-1,3-dioxane-4-carboxamide (S4a)

In a 10 mL pear-shaped flask, (4*R*)-*N*-(3-((2-aminoethyl)amino)-3-oxopropyl)-2-(4-methoxyphenyl)-5,5-dimethyl-1,3-dioxane-4-carboxamide **S3** (Meier & Burkart, 2009) (57.0 mg, 0.105 mmol, 1.0 equiv.), (*E*)-3-chloro-2-tetradecenoic acid **S2a** (39.2 mg, 0.1503 mmol, 1.0 equiv.), and 5 mL CH_2_Cl_2_ were added. To the solution was added PyBOP (94.3 mg, 0.181 mmol, 1.7 equiv.) and DIPEA (78.5 μL, 0.4509 mmol, 3.0 equiv.). After 20 hours, the volatiles were removed by rotary evaporation. Purification by silica flash chromatography (3:1 hexanes/ethyl acetate → 39:1 ethyl acetate/methanol) afforded **S4a** (64.8 mg, 69%, white solid).

**TLC:** R_f_ 0.31 (ethyl acetate). **^1^H-NMR** (400 MHz, CDCl3): δ 7.41 (d, *J* = 8.2 Hz, 2H), 7.02 (s, 1H), 6.89 (d, *J* = 8.3 Hz, 3H), 6.75 (s, 1H), 5.97 (s, 1H), 5.44 (s, 1H), 4.06 (s, 1H), 3.79 (s, 3H), 3.65 (q, *J* = 11.4 Hz, 2H), 3.50 (m, 2H), 3.31 (s, 3H), 2.97 (t, *J* = 7.0 Hz, 2H), 2.38 (bs, 2H), 1.58 (bs, 2H), 1.24 (s, 15H), 1.05 (d, *J* = 6.4 Hz, 6H), 0.84 (t, *J* = 6.3 Hz, 3H). **^13^C-NMR** (100 MHz, CDCl_3_): δ 172.05, 169.96, 164.95, 160.37, 153.67, 130.15, 127.66, 120.97, 113.83, 101.46, 83.89, 55.42, 39.84, 39.59, 36.36, 35.42, 35.05, 33.17, 32.01, 29.75, 29.73, 29.65, 29.54, 29.44, 29.05, 27.87, 22.78, 21.94, 19.21, 14.23.

#### (4*R*)-*N*-(3-((2-((*Z*)-3-chlorotetradec-2-enamido)ethyl)amino)-3-oxopropyl)-2-(4-methoxyphenyl)-5,5-dimethyl-1,3-dioxane-4-carboxamide (S4b)

Prepared as described for **S4a** from **S3** (Meier & Burkart, 2009) and **S2b** to afford **S4b** (116.3 mg, 85%, white solid). **TLC:** R_f_ 0.11 (ethyl acetate). **^1^H-NMR** (400 MHz, CDCl_3_): δ 7.39 (d, *J* = 8.7 Hz, 2H), 7.06 (t, *J* = 6.2 Hz, 1H), 6.95 (s, 2H), 6.87 (d, *J* = 8.7 Hz, 2H), 5.96 (s, 1H), 5.43 (s, 1H), 4.04 (s, 1H), 3.77 (s, 3H), 3.67 (m, 3H), 3.48 (m, 2H), 3.33 (s, 4H), 3.10 (m, 1H), 2.36 (m, 3H), 1.53 (s, 2H), 1.37 (q, *J* = 6.7, 6.1 Hz, 7H), 1.23 (s, 15H), 1.06 (d, *J* = 10.2 Hz, 6H), 0.83 (t, *J* = 6.6 Hz, 3H). **^13^C-NMR** (100 MHz, CDCl_3_): δ 172.19, 169.68, 165.22, 160.25, 144.34, 130.18, 127.61, 119.50, 113.75, 101.34, 83.84, 78.43, 55.36, 55.28, 43.24, 40.83, 39.53, 35.99, 35.05, 33.07, 31.93, 29.66, 29.64, 29.53, 29.37, 28.70, 27.28, 22.72, 21.86, 19.15, 18.56, 17.13, 14.16, 12.61.

#### (*R*,*E*)-3-chloro-*N*-(2-(3-(2,4-dihydroxy-3,3-dimethylbutanamido)propanamido)ethyl)tetradec-2-enamide ((*E*)-C14Cl)

In a 20 mL vial, **S4a** (64.8 mg, 0.104 mmol, 1.0 equiv.) and 1.0 mL 4:1 AcOH/H_2_O were added. After 21 hours, the mixture was concentrated by rotary evaporation, and azeotroped from cyclohexane (3×10 mL) and benzene (3×10 mL). Purification by silica flash chromatography (ethyl acetate → 85:15 ethyl acetate/methanol) afforded **((*E*)-C14Cl** (38.2 mg, 73%, clear oil).

**TLC:** Rf 0.31 (9:1 ethyl acetate/methanol). **^1^H-NMR** (400 MHz, CDCl_3_): δ 7.54 (t, *J* = 5.1 Hz, 1H), 7.30 (s, 1H), 7.17 (s, 1H), 6.03 (s, 1H), 3.98 (s, 1H), 3.68–3.21 (m, 9H), 2.95 (t, 2H), 2.43 (s, 2H), 1.58 (s, 2H), 1.24 (s, 18H), 0.89 (d, *J* = 14.4 Hz, 6H), 0.84 (t, *J* = 6.6 Hz, 3H). **^13^C-NMR** (100 MHz, CDCl_3_): δ 174.38, 172.55, 165.33, 153.90, 120.96, 77.58, 77.16, 76.74, 70.73, 39.86, 39.49, 39.37, 36.10, 35.50, 32.04, 29.79, 29.76, 29.68, 29.58, 29.47, 29.12, 27.92, 22.81, 21.34, 20.82, 14.25.

#### (*R*,*Z*)-3-chloro-*N*-(2-(3-(2,4-dihydroxy-3,3-dimethylbutanamido)propanamido)ethyl)tetradec-2-enamide ((*Z*)-C14Cl)

Prepared as described for **(*E*)-C14Cl** from **S4b** to afford **((*Z*)-C14Cl** (35.9 mg, 38%, white solid).

**TLC:** R_f_ 0.29 (9:1 ethyl acetate/methanol). **^1^H-NMR** (400 MHz, CDCl_3_): **^13^C-NMR** (100 MHz, CDCl_3_):

### C. NMR Spectra

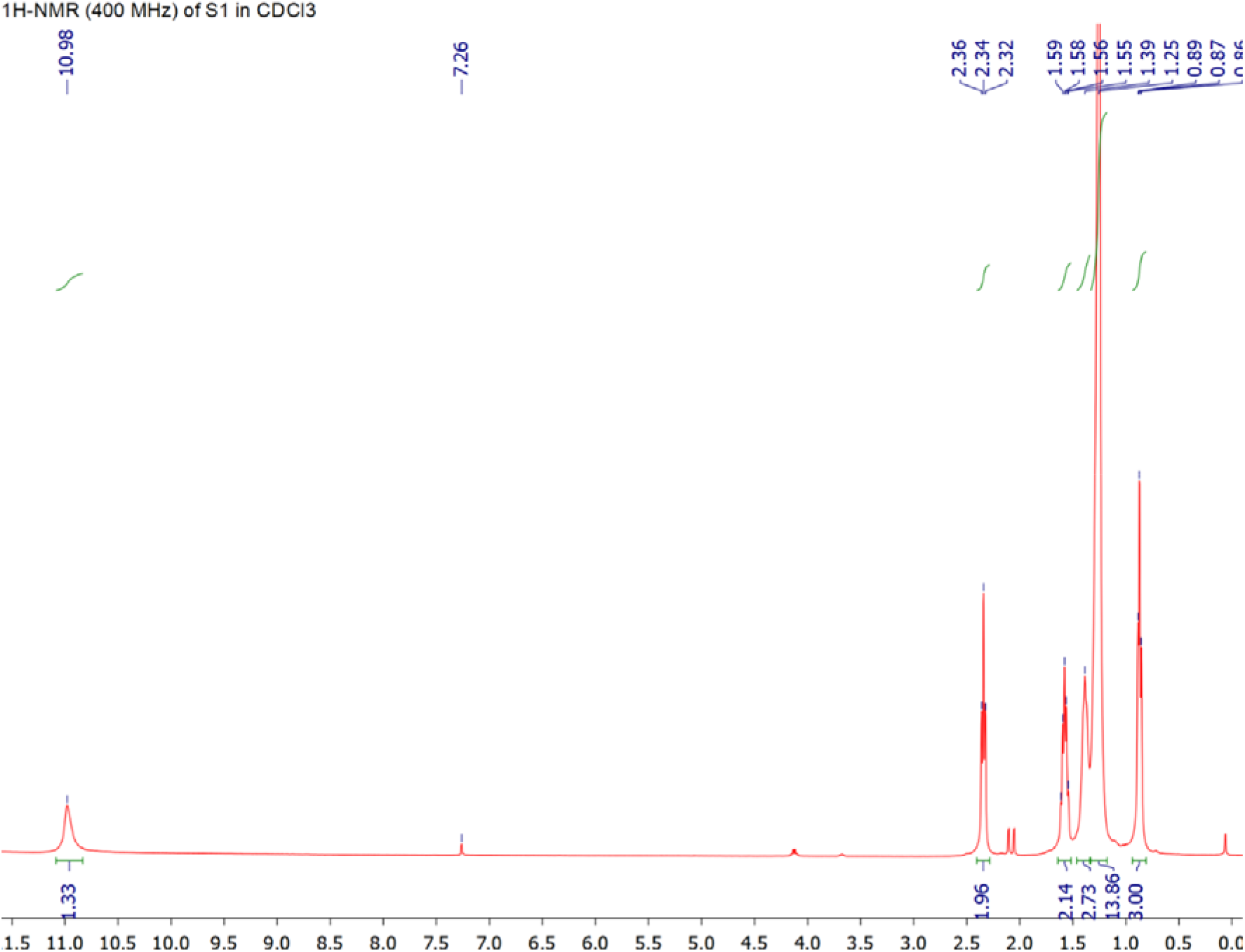

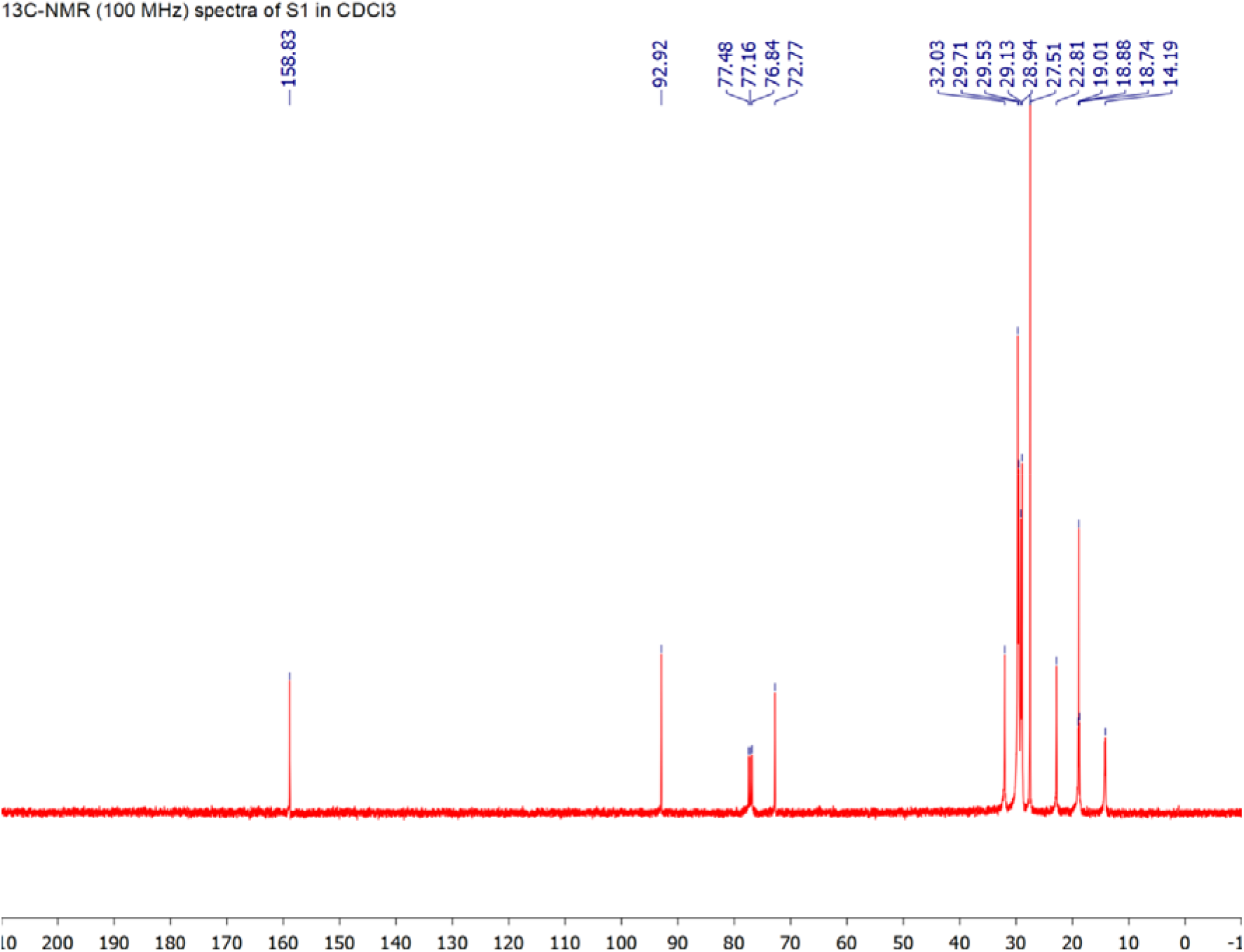

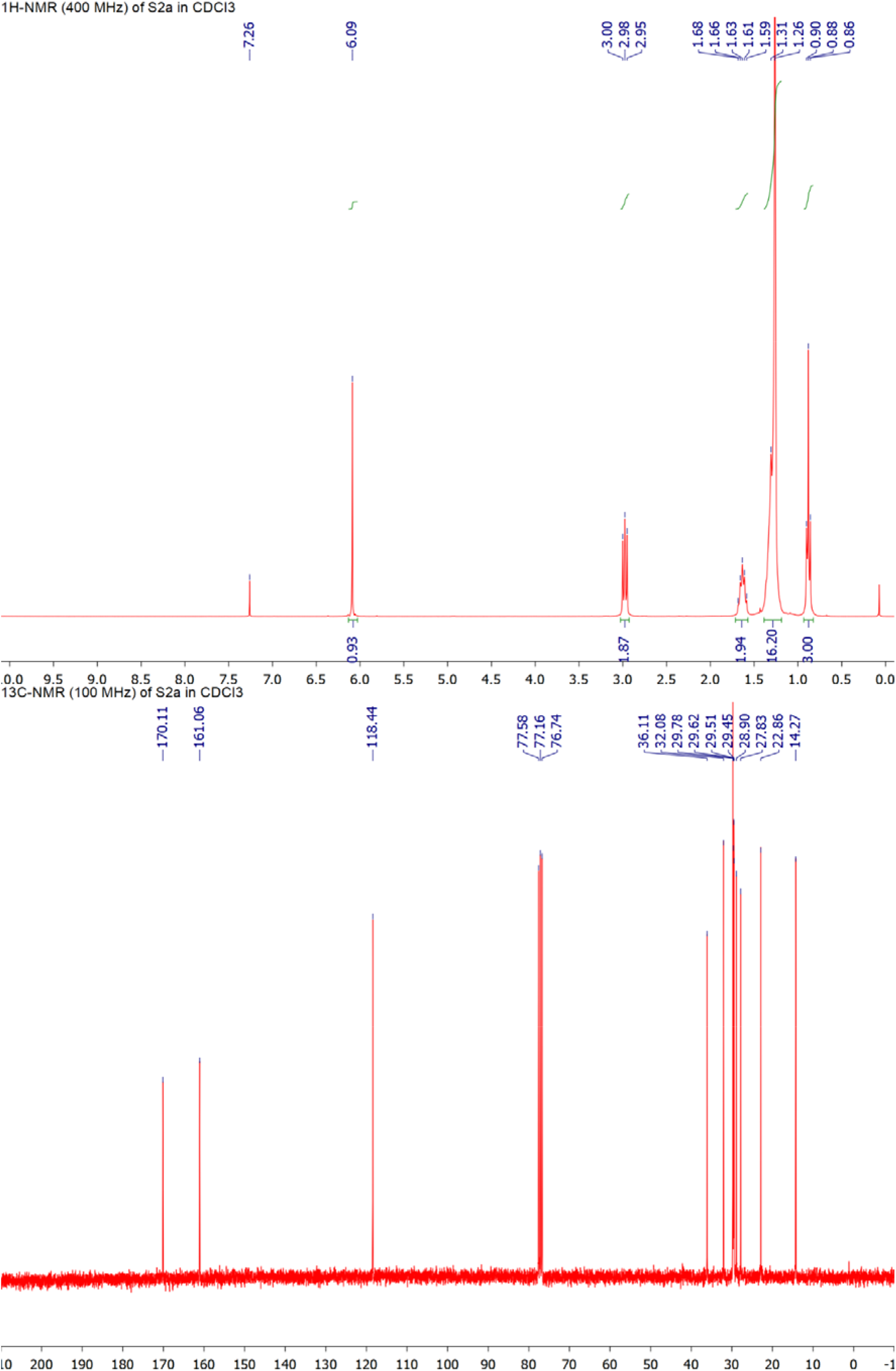

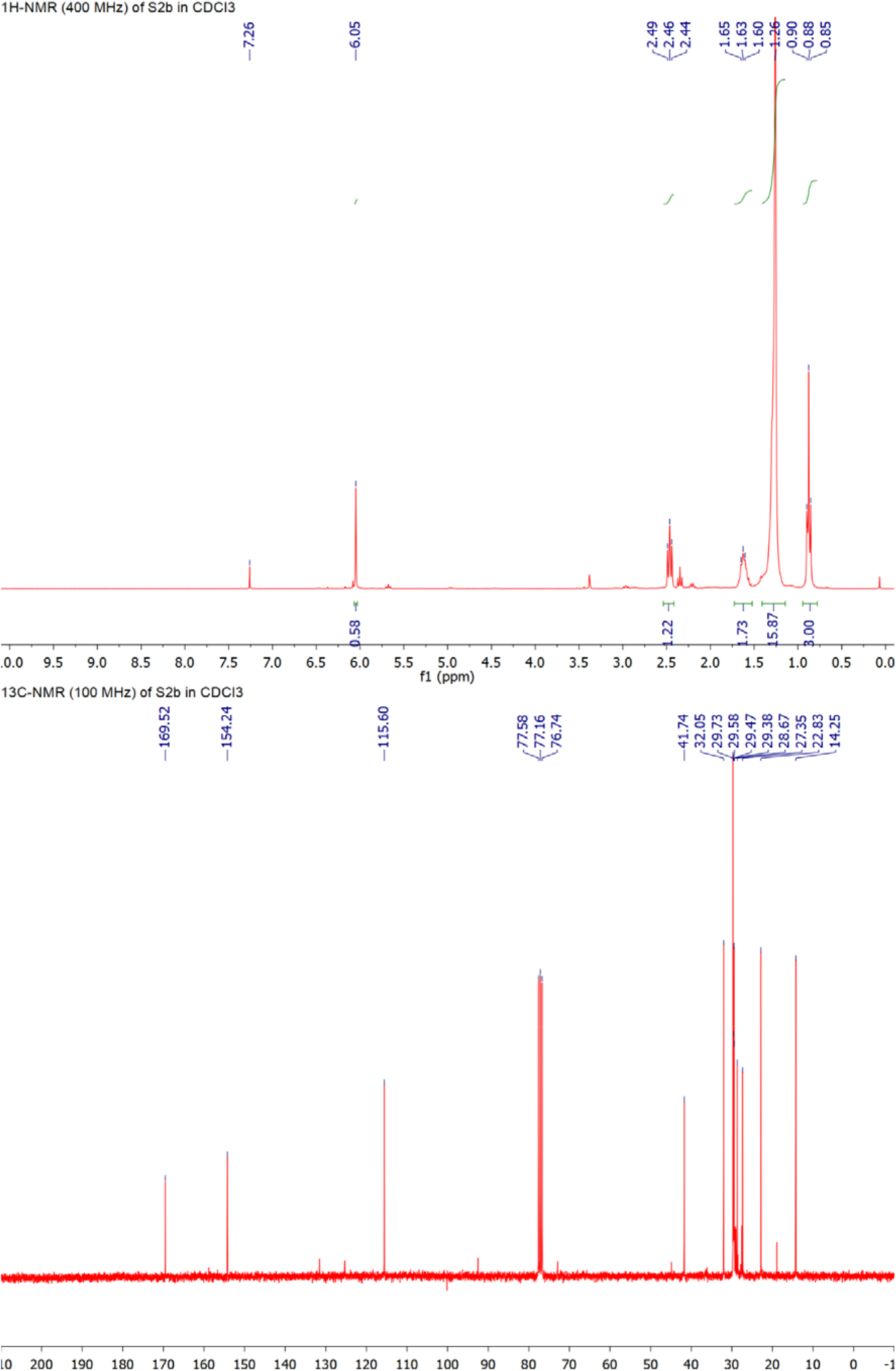

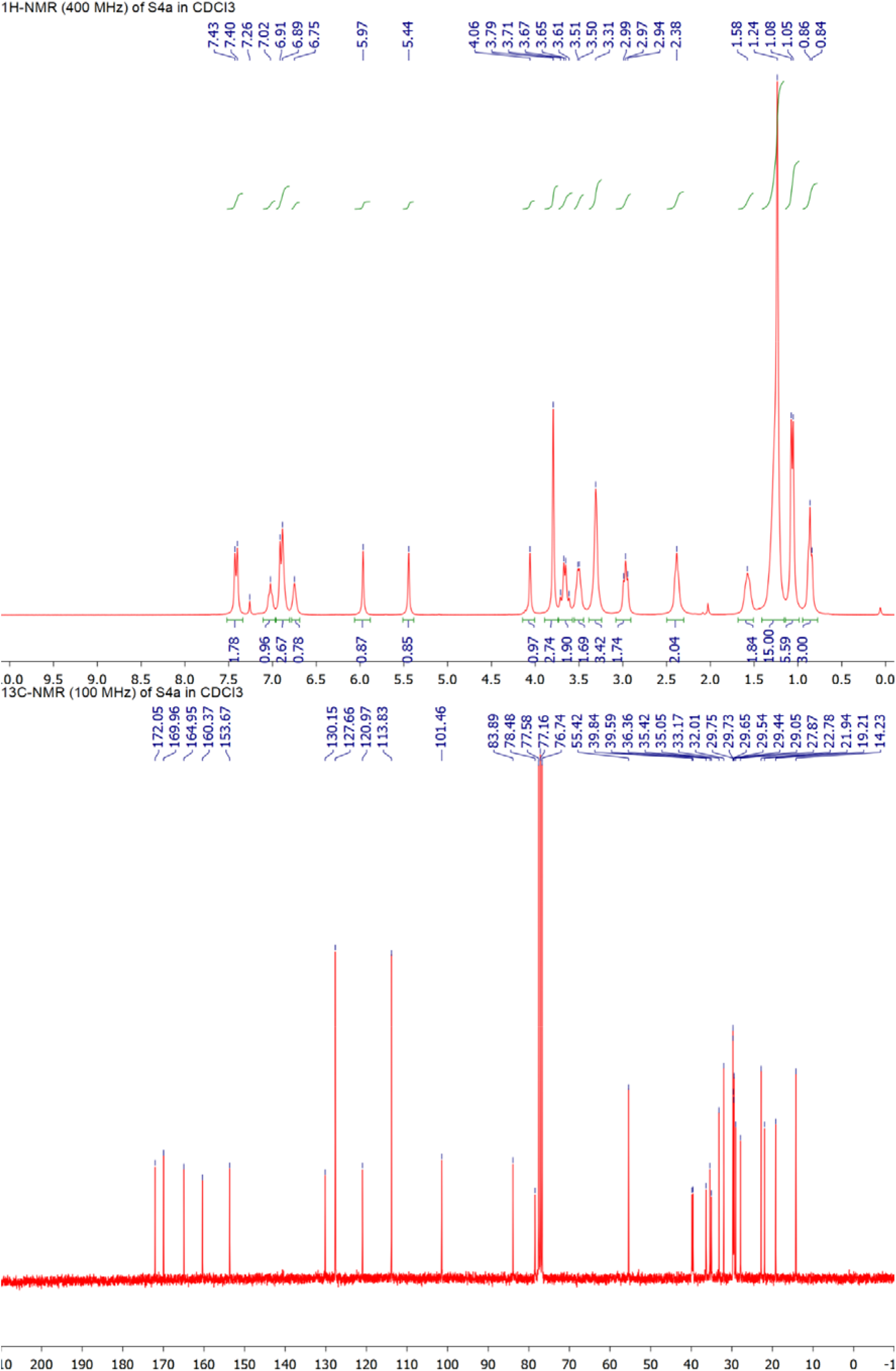

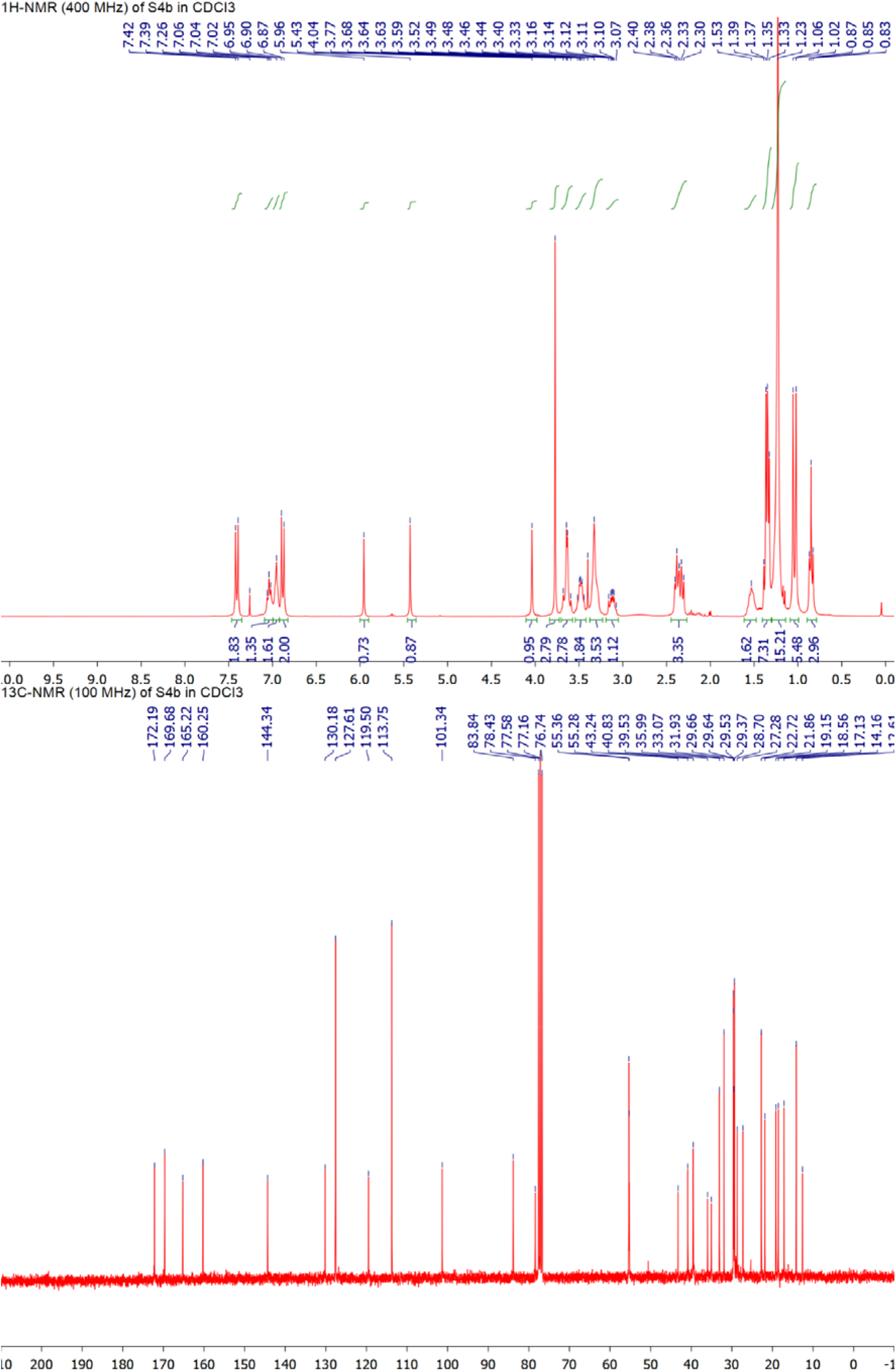

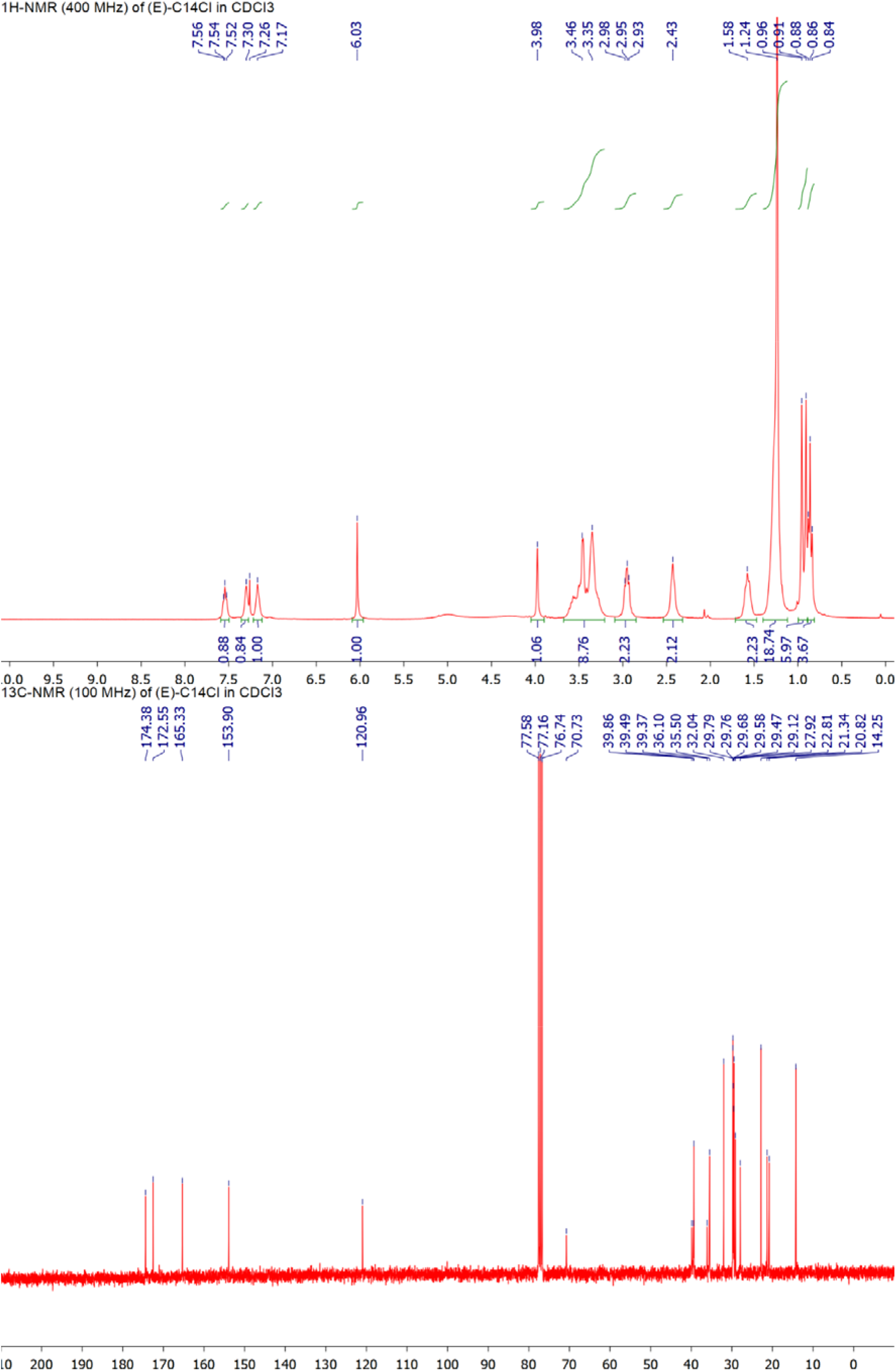

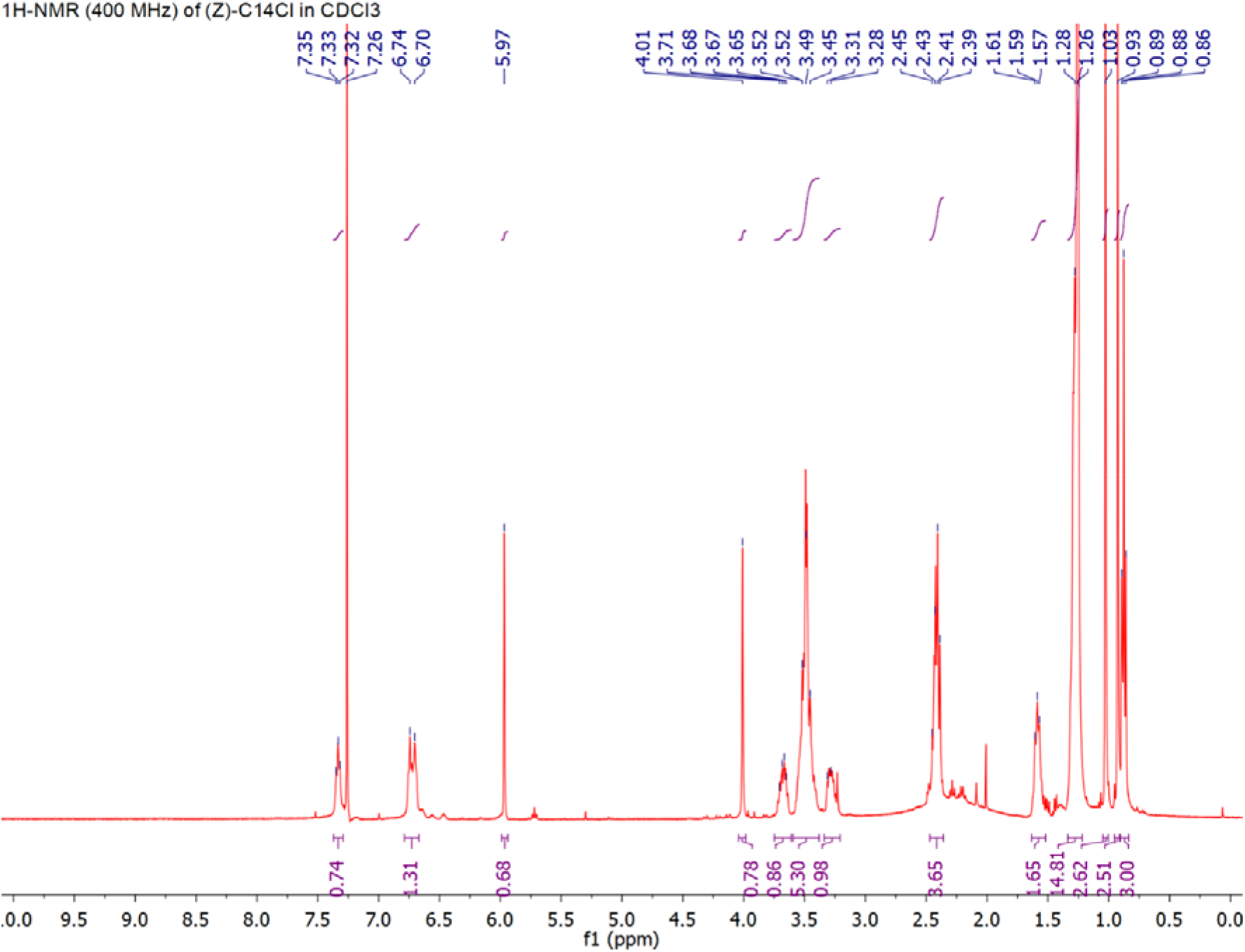

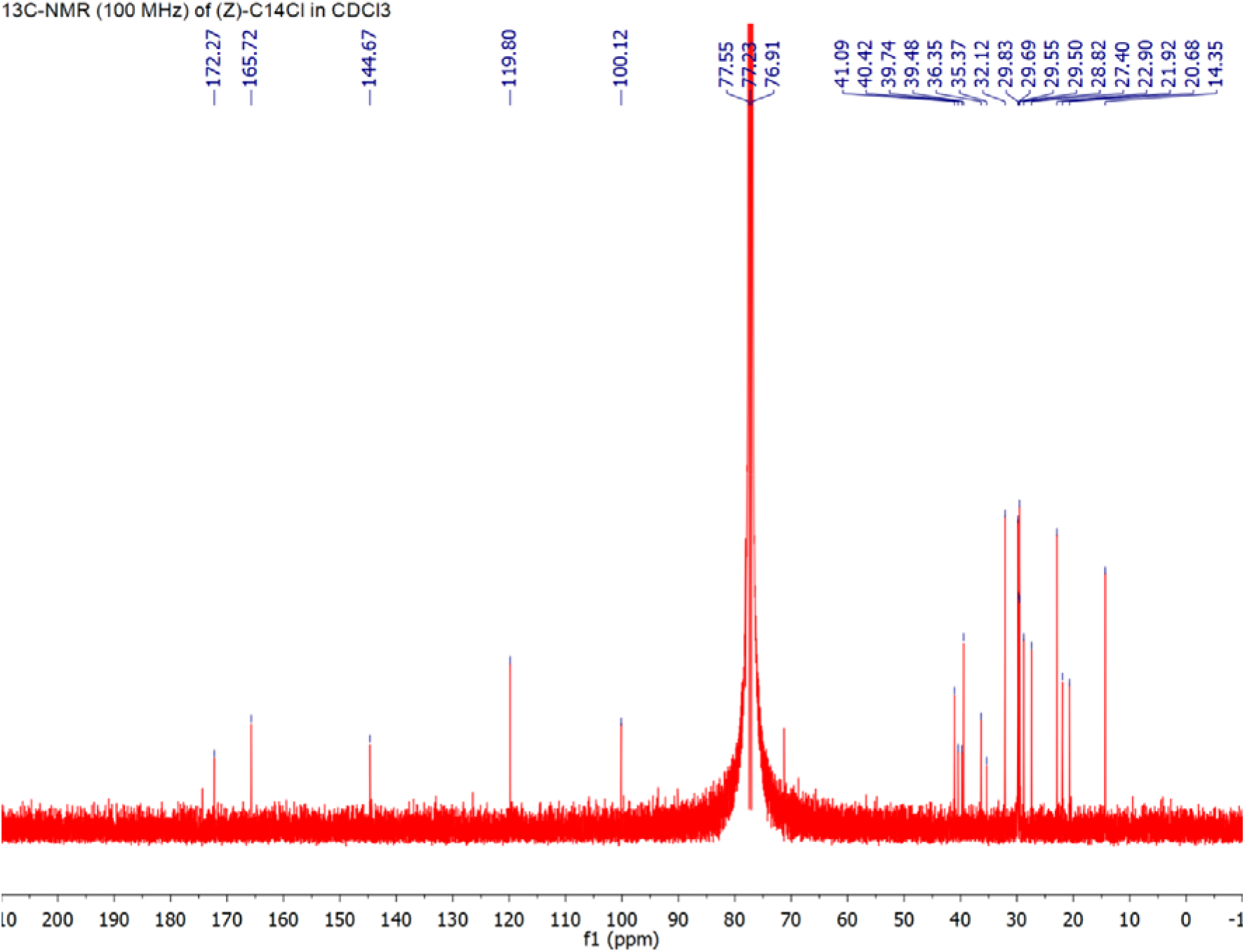

## Notes

### Competing Interest Statement

The authors have declared no competing interest.

